# Vascular and perivascular cell profiling reveals the molecular and cellular bases of blood-brain barrier heterogeneity

**DOI:** 10.1101/2021.04.26.441465

**Authors:** Sarah J. Pfau, Urs H. Langen, Theodore M. Fisher, Indumathi Prakash, Faheem Nagpurwala, Ricardo A. Lozoya, Wei-Chung Allen Lee, Zhuhao Wu, Chenghua Gu

**Affiliations:** Department of Neurobiology, Harvard Medical School, Boston, MA 02115, USA; Department of Cell, Developmental & Regenerative Biology and Department of Neuroscience, Icahn School of Medicine at Mount Sinai, New York, NY 10029, USA

## Abstract

SUMMARY

The blood-brain barrier (BBB) is critical for protecting the brain and maintaining neuronal homeostasis. Although the BBB is a unique feature of the central nervous system (CNS) vasculature, not all brain regions have the same degree of impermeability. Differences in BBB permeability are important for controlling the local extracellular environment of specific brain regions to regulate the function and plasticity of particular neural circuits. However, how BBB heterogeneity occurs is poorly understood. Here, we demonstrate how regional specialization of the BBB is achieved. With unbiased cell profiling in small, defined brain regions, we compare the median eminence, which has a naturally leaky BBB, with the cortex, which has an impermeable BBB. We identify hundreds of molecular differences in endothelial cells (ECs) and demonstrate the existence of differences in perivascular astrocytes and pericytes in these regions, finding 3 previously unknown subtypes of astrocytes and several key differences in pericytes. By serial electron microscopy reconstruction and a novel, aqueous-based tissue clearing imaging method, we further reveal previously unknown anatomical specializations of these perivascular cells and their unique physical interactions with neighboring ECs. Finally, we identify ligand-receptor pairs between ECs and perivascular cells that may regulate regional BBB integrity in ECs. Using a bioinformatic approach we identified 26 and 26 ligand-receptor pairs underlying EC-pericyte and EC-astrocyte interactions, respectively. Our results demonstrate that differences in ECs, together with region-specific physical and molecular interactions with local perivascular cells, contribute to BBB functional heterogeneity. These regional cell inventories serve as a platform for further investigation of the dynamic and heterogeneous nature of the BBB in other brain regions. Identification of local BBB specializations provides insight into the function of different brain regions and will permit the development of region-specific drug delivery in the CNS.

## INTRODUCTION

The BBB is a physiological barrier between the blood and brain that maintains the ionic and chemical composition of the brain extracellular environment for proper neuronal function. BBB breakdown is involved in neurodegenerative diseases^1^. Conversely, an intact BBB is a major obstacle for drug delivery to the CNS to treat neurological disorders^2^. Understanding the molecular mechanisms of BBB regulation will permit manipulation of the barrier in either direction to promote disease treatment or barrier repair.

Although the BBB is a unique feature of the CNS vasculature, not all brain regions have the same degree of impermeability. Seven specialized regions known as the circumventricular organs (CVOs) have a naturally leaky BBB, which is vital for their functions. Neurons in CVOs sense signaling compounds and secrete brain-derived hormones in the systemic circulation, facilitating rapid communication between the brain and the periphery to regulate feeding behaviors, cardiovascular function and thirst^3, 4^. BBB heterogeneity has also been observed in the hippocampus, basal ganglia and cerebellum, where increased BBB permeability was observed in human aging and with the early onset of neurodegenerative diseases^1, 5, 6^. Yet, how these naturally existing variations in BBB permeability occur is poorly understood.

CNS capillary endothelial cells (cECs) constitute the BBB and restrict access between the blood and brain with specialized tight junctions between cells and a low rate of transcytosis^1, 7–9^. Thus, previous work has focused largely on identifying key molecular determinants of the BBB in CNS ECs by bulk RNA sequencing, comparing ECs from the brain with peripheral tissues^10–13^. However, BBB properties are not entirely intrinsic to ECs but require active induction and maintenance from other perivascular cells^14–18^. Specifically, pericytes and astrocyte endfeet ensheath brain capillaries, forming the interface between ECs and neurons. Remarkably, each neuron is within 15 microns of a capillary^19^, suggesting that these interactions are important for the dynamic regulation of BBB permeability. Indeed, mice with reduced numbers of pericytes and astrocytes have a leaky BBB^16–18, 20^, and major facilitator superfamily domain containing 2A (*Mfsd2a*), a key BBB regulator in brain cECs, is downregulated in brain ECs in pericyte-deficient mice^10^. Currently, how intercellular communication between vascular and perivascular cells and their local organization contributes to regional differences in BBB permeability is poorly understood.

The major technical challenge to uncovering the mechanism underlying BBB heterogeneity is that ECs are rare in the brain compared to other cell types, representing around 5% of brain cells^11, 21^. Thus, while typical unbiased single-cell gene expression studies of the brain often include vascular cells, they yield limited data about their transcriptomes due to their relative scarcity following dissociation protocols optimized for neurons^22–24^. To circumvent this problem, most studies of brain vascular and perivascular cells have relied on cell sorting from the entire brain or from large brain regions^13, 25, 26^. This pooling-based approach fails to capture BBB heterogeneity because it underrepresents smaller brain regions. Such regions may contain transcriptionally diverse and specialized cells. Therefore, to go beyond population-level characterization of brain vascular and perivascular cells to investigate regional heterogeneity necessitates the development of methods to enrich for ECs and to discern regional differences in BBB-associated cells, especially in small brain regions such as the CVOs.

Here, we develop a platform to comprehensively understand how vascular and perivascular cells affect BBB functional heterogeneity. We perform unbiased, high-throughput single-cell RNA sequencing (scRNAseq) of one of the CVOs, the median eminence (ME), and a size-matched region (∼0.05 × 0.2 × 1.2 mm^3^) of the somatosensory cortex (cortex). Comparison of these two small brain regions with distinct BBB properties revealed significant molecular differences in cECs and perivascular astrocytes. Consistent with this molecular heterogeneity, we observed significant morphological heterogeneity in vascular and perivascular cells in these two regions by electron microscopy and three-dimensional whole brain imaging following the newly-developed U.Clear tissue clearing method (Farrell R., Wang W. and Wu Z, *in submission*). Moreover, these morphological differences were accompanied by striking differences in the local organization of vascular and perivascular cells. Finally, bioinformatics analyses revealed key pericyte- and astrocyte-derived signals that may act upon ECs to confer barrier properties. Taken together, this work reveals how physical interactions and communication between cECs and neighboring pericytes and astrocytes contributes to BBB functional heterogeneity within the brain. These findings highlight the importance of considering regional vascular and perivascular cell diversity in understanding BBB heterogeneity and developing region-specific therapies.

## RESULTS

### Morphological, molecular and functional differences of the brain vasculature between the ME and cortex visualized with U.Clear

To characterize the vasculature in the ME and cortex, we employed U.Clear (Farrell R., Wang W. and Wu Z, in submission). U.Clear is a novel, aqueous-based tissue clearing protocol that preserves endogenous fluorescence and permits the use of most antibodies to stain intact, adult mouse tissues in their entirety. U.Clear provided us with an unprecedented, three-dimensional view of the entire, intact ME and a size-matched region of the cortex after confocal microscopy imaging, while still providing fine molecular and morphological details at higher magnifications. As shown previously^27–30^, the entire ME vascular network leaks tracer into the ME while the entire vascular network in the cortex region does not permit passage of tracer into the cortex (Fig. 1a, Extended Data Fig. 1a, Supplementary Video 1). Moreover, cortical capillaries form a network of thin, evenly spaced vessels while the ME vasculature is a more densely arranged plexus of wider, tortuous vessels (Fig. 1a-b, Supplementary Video 2). We find larger capillary diameter in the ME than in the cortex (Fig. 1c) and increased EC density in ME capillaries when compared to the cortex (Fig. 1d, Extended Data Fig. 1b). Finally, distinct molecular expression was observed in ECs in each of these two regions. ME ECs express Plasmalemma vesicle-associated protein (Plvap), a component of EC stomatal and fenestral diaphragms that is also expressed in peripheral ECs^31, 32^. Plvap is completely absent in cortical ECs (Fig. 1e, Extended Data Fig. 1c). In contrast, cortical ECs express glucose transporter 1 (Glut1, encoded by the gene *Slc2a1*), Claudin-5 (Cldn5) and Mfsd2a throughout the capillary network while both are completely absent in ME ECs (Extended Data Fig. 1c-e). Thus, we characterized the vasculature in each region in its entirety with U.Clear, observing clear differences in BBB permeability and vessel morphology together with molecular differences in cECs.

**Fig. 1.**
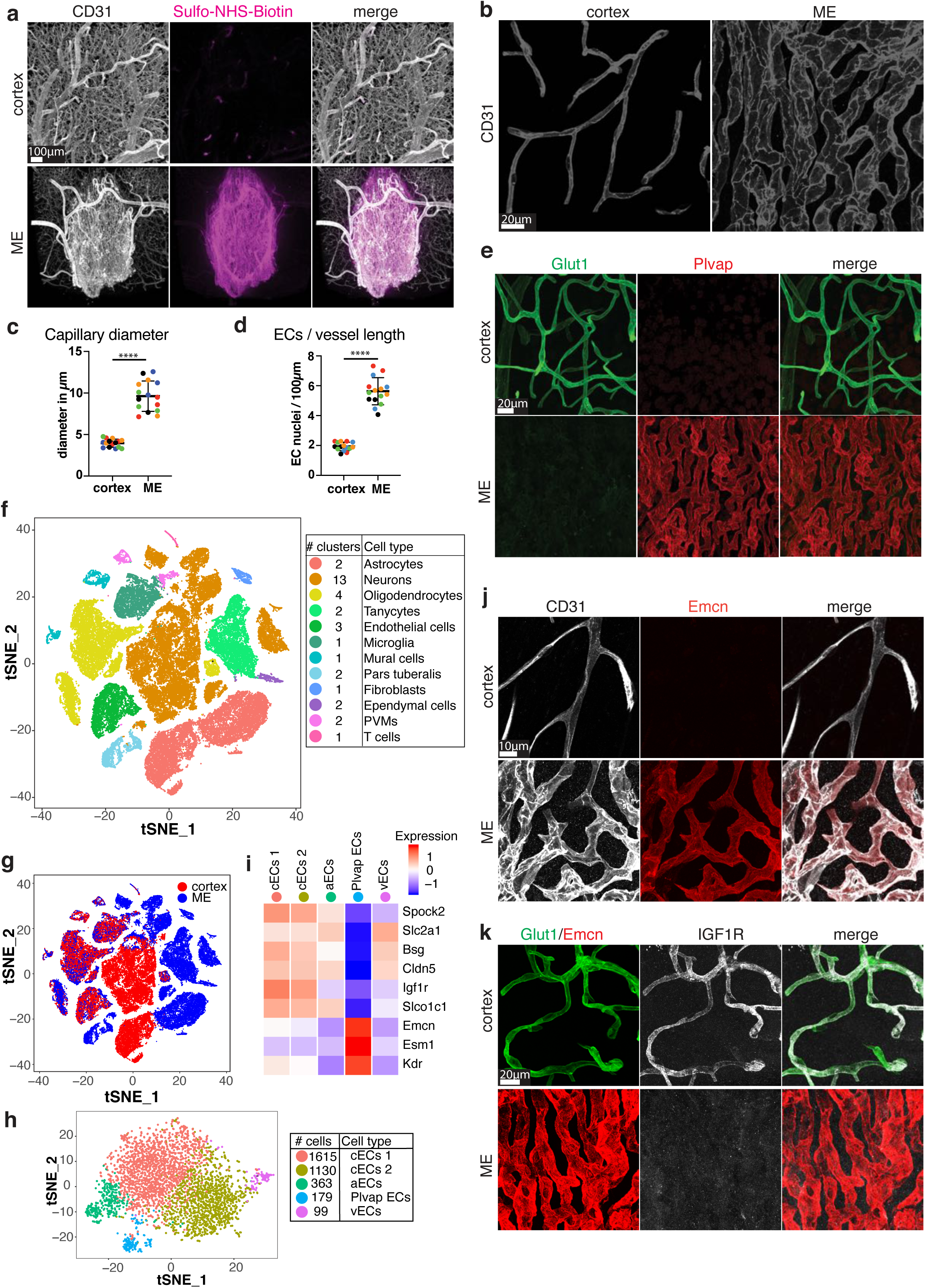
Morphological, molecular and functional differences of the vasculature between the ME and cortex. (a) Tracer leakage assay using Sulfo-NHS-Biotin (magenta) as tracer and immunostaining for blood vessels (CD31, white) in cortex (upper panel) and ME (lower panel). Tracer in circulation has been washed out by perfusion prior to analysis. Scale bar 100µm. (b) High magnification images of capillaries (CD31) highlighting vessel morphology in cortex (left) and ME (right). Scale bar 20µm. (c) Quantification of capillary diameter in cortex and ME. (n=5 mice, for quantification 3 images per region and mouse were taken, same colors refer to same mice, data presented as mean±SD, p<0.0001, nested two-tailed t-test). (d) Quantification of endothelial cell nuclei (ERG+) over length of capillaries. (n=5 mice, for quantification 3 images per region and mouse were taken, same colors refer to same mice, data presented as mean±SD, p<0.0001, nested two-tailed t-test). (e) High magnification images of capillaries showing distinct Glut1 (green) and Plvap (red) expression pattern and vessel morphology in the cortex and ME. Scale bar 20µm. (f) tSNE projection of 65,872 single-cell transcriptomes (32,183 from ME and 33,689 from cortex). Cell type clusters were color-coded and annotated post hoc based on their transcriptional profiles (see details in Methods). The number of clusters identified for each cell type is indicated in the plot legend. (g) tSNE projection of the dataset in (f) color coded by sample region. (h) tSNE projection of 3,933 EC transcriptomes. 5 subtypes of ECs were identified with an unbiased analysis and cell types were annotated based on their transcriptional profiles (see supplemental figure). The number of each cell type profiled is indicated in the plot legend. (i) Heatmap illustrating the average relative expression of region-specific genes in each subtype cluster identified in (h). (j) Co-immunostaining of Emcn (red) with CD31 in cortex and ME, validating that Emcn is ME cEC enriched. Scale bar 10µm. (k) Co-immunostaining for cortex cEC-enriched protein Igf1R (white), ME cEC-enriched protein Emcn (red), and cortex cEC-enriched protein Glut1 (green) in cortex and ME. Scale bar 20µm.

### scRNAseq of ME and a size-matched region of cortex reveals unique cell types and subtypes in each brain region

To unbiasedly reveal molecular differences between these two regions, we next performed inDrops scRNAseq^33, 34^ of mouse cells from the ME and a size-matched region of the cortex (Extended Data Fig. 2a). We developed a tissue dissociation protocol to obtain efficient, unbiased recovery of vascular cells. All cell types associated with brain blood vessels are well-represented in our dataset, and ECs and mural cells comprise ∼5% of the cells analyzed, on par with estimates of the prevalence of these cell types in the mouse brain^11, 21^. Unbiased cell clustering with Seurat identified cell type-specific clusters (Fig. 1f)^35, 36^, and we analyzed ME and cortex samples together to perform direct comparisons. After quality control filtering (see Methods), 65,872 high quality cells were retained for further analysis, 32,183 from the ME and 33,689 from the cortex. We identified 34 clusters that correspond to 12 major cell types based on expression of cell-type-specific transcripts (Fig. 1f, Extended Data Fig. 2b, 3a-e, Table 1): neurons, oligodendrocytes, astrocytes, tanycytes, ECs, microglia, mural cells, pars tuberalis cells, perivascular macrophages (PVMs), fibroblasts, ependymal cells and T cells. Interestingly, an ME-specific EC cluster (ECs 2) was observed (Fig. 1g, Extended Data Fig. 2c). ECs also comprised two other clusters, which are derived from the cortex and, in the case of *Plvap*-negative ME ECs, likely from regions adjacent to the ME that contain blood vessels with barrier properties that are inevitably included in our dissection. Moreover, astrocytes from the ME and cortex clustered separately (Fig. 1g, Extended Data Fig. 2c). Mural cells from both regions were found in the same cluster (Fig. 1g, Extended Data Fig. 2c).

**Fig. 2.**
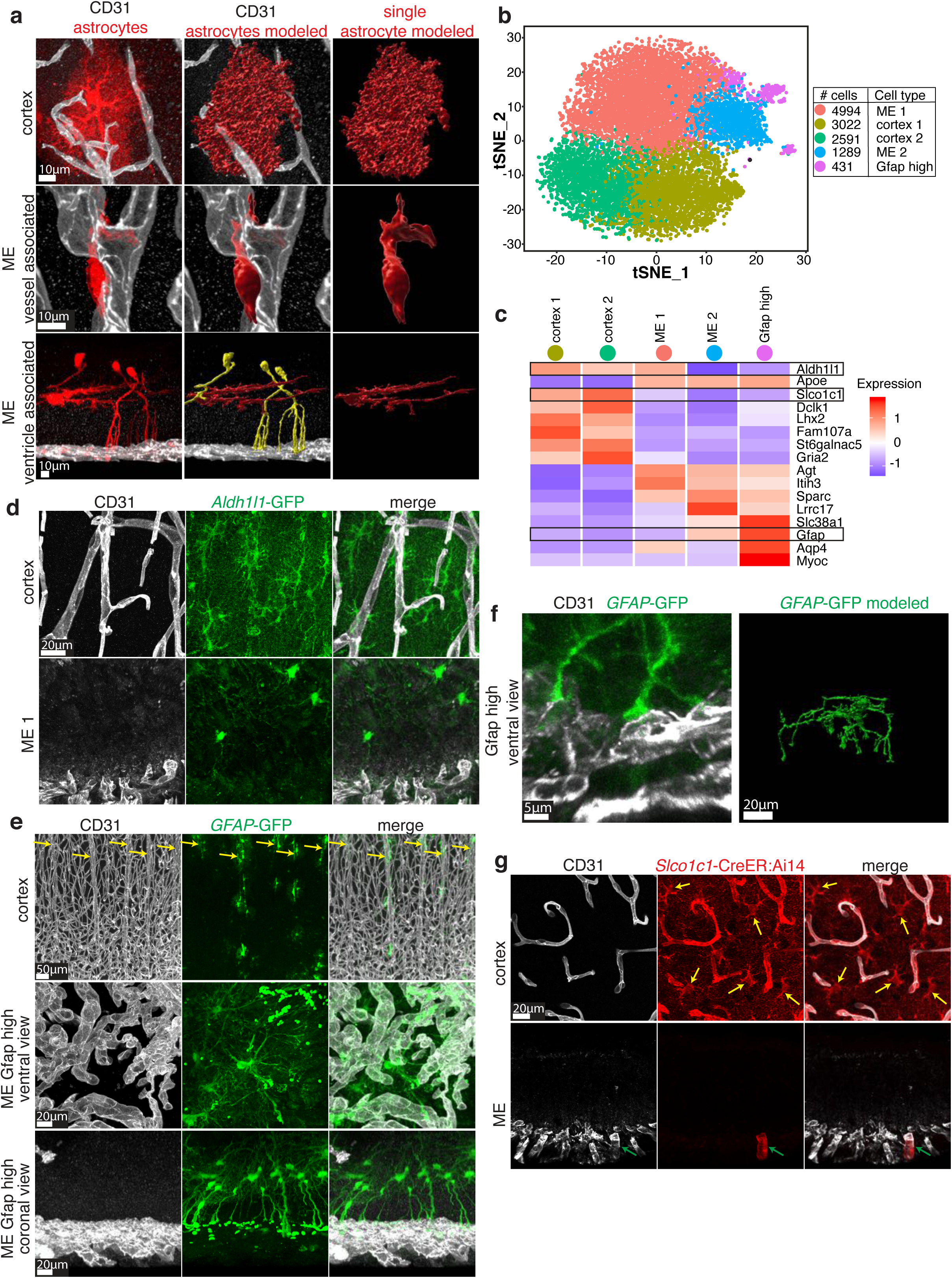
Distinct astrocyte subtypes with unique interactions with blood vessels found between the ME and cortex. (a) Fluorescent labeling of astrocyte populations in cortex and ME using Glast-CreER:Ai14 mice after low dose 4OH-tamoxifen to achieve sparse cell labeling. Left column: immunostaining for Tomato-positive astrocytes (red) and blood vessels (CD31, white). Middle and right column display 3D reconstructions of astrocytes (red). Cells modeled in yellow are tanycytes. Scale bar 10µm. For a comparison of different astrocyte populations with same scales see Extended Data Fig. 5b. (b) tSNE projection of 12,327 astrocyte transcriptomes. Astrocyte subtype clusters were identified with an unbiased analysis. The number of each cell subtype profiled is indicated in the plot legend. (c) Heatmap illustrating the average relative expression of differentially expressed genes in each cluster identified in (b). (d) Fluorescent labeling (green) of cortex astrocytes and ventricle-associated ME 1 astrocytes using *Aldh1l1*-GFP mice. Co-staining for CD31 (white) to label capillaries. Scale bar 20µm. (e) Co-staining for CD31 (white, vessels) and GFP in the cortex and ME of GFAP-GFP mice. Note that GFP labels ventricle-associated Gfap high ME astrocytes shown both in ventral and coronal view. In cortex GFP only sparsely labeled peri-arterial astrocytes (yellow arrows). Scale bar 50µm in top row and 20µm in middle and lower row. (f) Left panel: High magnification of GFAP-high ME astrocyte interaction with capillaries shown in (f). Scale bar 5µm. Right panel: Imaris 3D-reconstruction of Gfap-high ventricle-associated ME astrocyte. Scale bar 20µm. (g) Co-staining of CD31 (blood vessels, white) with Tomato in the cortex and ME of *Slco1c1*- CreER:Ai14 mice. Tomato (red) indicates Slco1c1 expression. Note Tomato expression in cortex capillaries and astrocytes, but not ME capillaries and astrocytes. Single Tomato positive vessel in ME (green arrow) is arterial. Yellow arrows point at astrocytes in cortex. Scale bar 20µm.

**Fig. 3.**
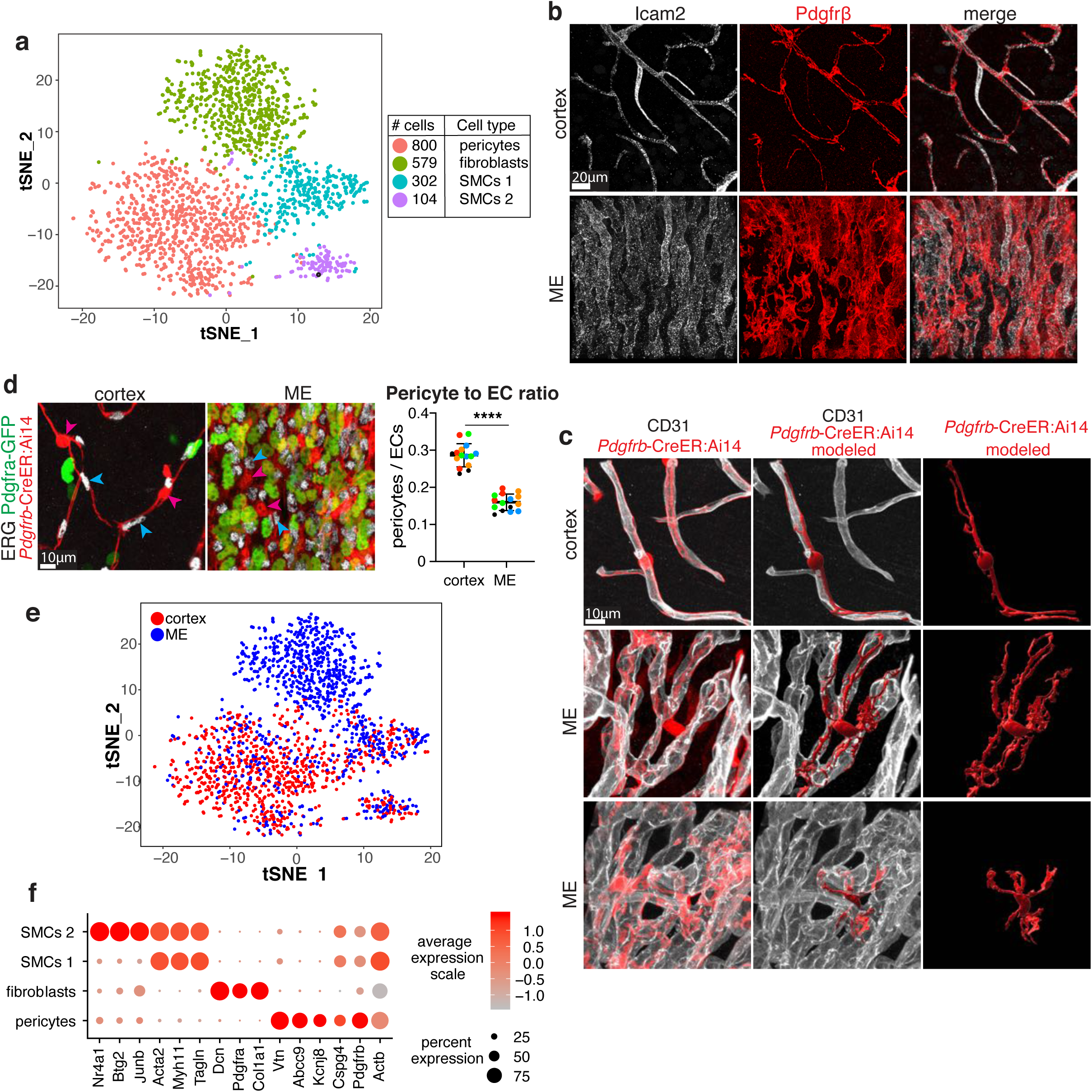
Pericytes associated with cortex and ME blood vessels show distinct molecular, morphological and anatomical features. (a) tSNE projection of 1,785 mural cell transcriptomes. Cell clusters were identified with an unbiased analysis. The number of each cell type profiled is indicated in the plot legend. Cell types were annotated based on their expression of cell type-specific gene transcriptional profiles (f). (b) Co-immunostaining for pan-pericyte marker Pdgfrβ (red) and pan-EC marker Icam2 (white) in cortex and ME. Scale bar 20µm. (c) Immunostaining and 3D-reconstruction using Imaris of single Tomato positive pericytes (red) in touch with capillaries (CD31, white) in cortex and ME (two examples). Single pericytes labelled by single low dose injection of 4OH-tamoxifen in adult *Pdgfrb*-CreER:Ai14 mice one week prior to analysis. Scale bar 10µm. (d) Left: Co-immunostaining for EC nuclei marker ERG (white) and pericytes labelled using *Pdgfrb*-CreER:Ai14 *Pdgfra*-GFP mice. Cyan arrowheads point at GFP-, Tomato+ pericytes, magenta arrowheads point at ERG+ EC nuclei. Scale bar 10µm. Right: Quantification of pericyte (GFP-, Tomato+) to EC (ERG+) ratio using *Pdgfra*-H2B-GFP;*Pdgfrb*-CreER:Ai14 mice (n=5 mice, for quantification 3 images per region and mouse were taken, same colors refer to same mice, data presented as mean±SD, p<0.0001, nested two-tailed t-test). (e) tSNE projection in (a) colored by sample region. (f) Dot plot of average expression of general mural cell markers and cell type-specific transcripts used to annotate cluster cell types in (a).

**Table 1.** scRNAseq cell type- and subtype-enriched genes and ligand-receptor interaction scores.

EC subclustering analysis uncovered 5 EC subtypes, including arteriolar ECs (aECs), cECs (cECs 1 and 2), and venous ECs (vECs) (Fig. 1h). Importantly, an ME-specific EC subcluster characterized by expression of *Plvap* corresponds to the rare, ME-specific EC population we observed by immunostaining (Extended Data Fig. 4a-b, Table 1). We further determined that *Plvap*-expressing ECs are capillary ECs (cECs) based on their lack of expression of arteriolar and venous markers (Extended Data Fig. 4b). In all analyses below, *Plvap*-positive cECs were compared with cortical cECs. Thus, our dissociation method captures rare EC subtypes in small brain regions as well as ECs from all segments of the vascular tree, permitting fine-grained molecular analysis of all cells in these regions and demonstrating that we can effectively investigate EC and perivascular cell heterogeneity with our method.

**Fig. 4.**
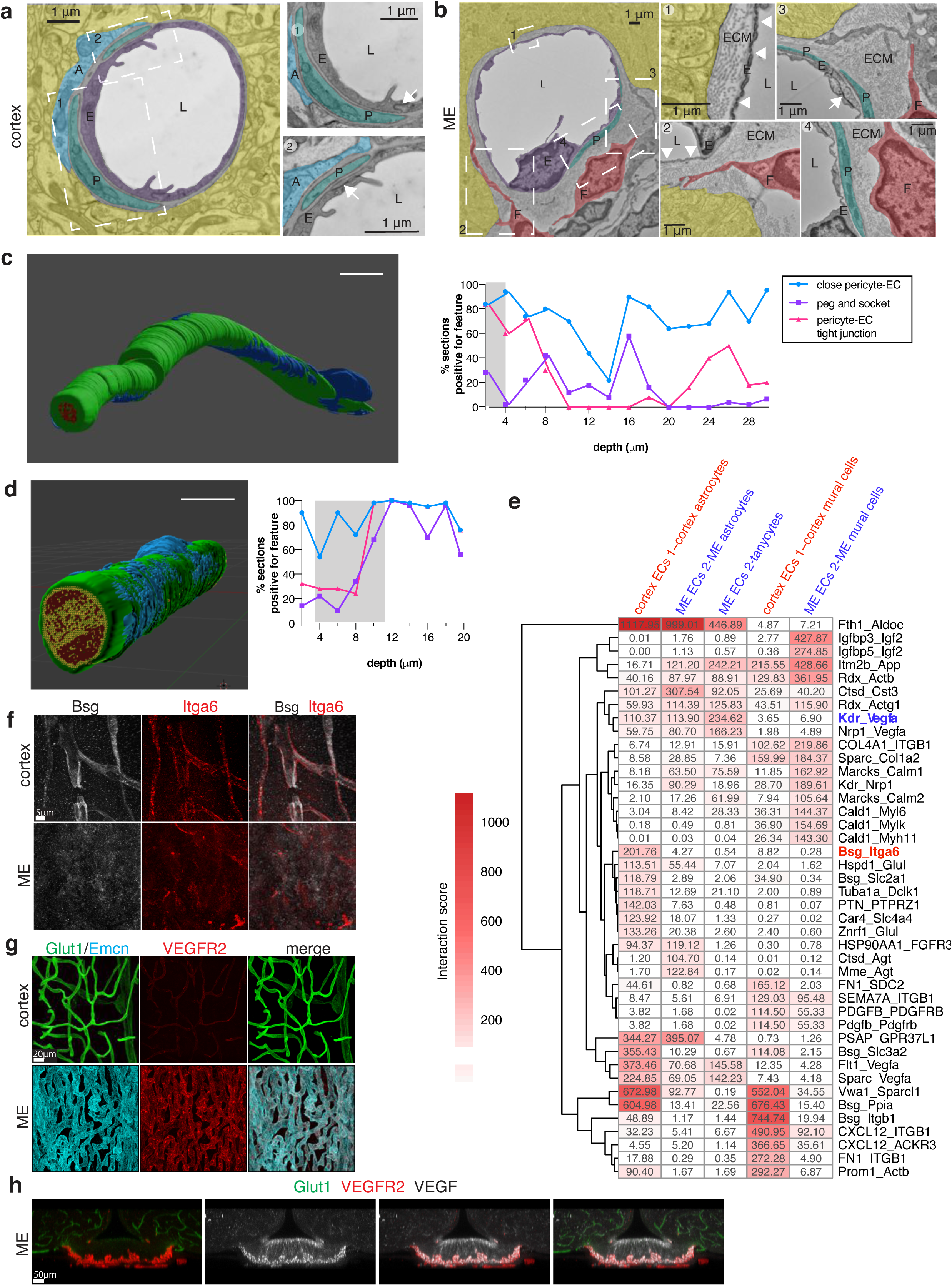
Imaging and bioinformatics analyses reveal differences in the organization of the vascular and perivascular cell environment and intercellular signaling capacity between the ME and cortex. (a) TEM images of cortical capillaries. Pseudocolors highlight different cells: cEC (E, purple), pericyte (P, teal), astrocyte endfoot (A, cyan), lumen (L, white) and neuropil (yellow). Insets show cEC tight junctions (white arrows), pericyte cells (P, teal) and astrocyte endfeet (A, cyan). Scale bar 1µm. (b) TEM images of a blood vessel in the ME. Pseudocolors highlight different cells: cEC (E, purple), pericyte (P, teal), fibroblast (F, red), lumen (L, white) and neuropil (yellow). Insets show capillary fenestrations (white arrowheads), cEC tight junctions (white arrows), extracellular matrix-filled perivascular space (ECM), pericyte cells (P, teal) and fibroblast cells (F, red). Scale bar 1 µm. (c) and (d) Serial TEM reconstruction of EC and pericyte interactions in two cortical blood vessels with quantification of three features of pericyte-EC interaction. In the reconstruction, ECs are shown in green, pericyte cells in blue and the blood vessel lumen in red. Scale bar represents 5 µm. Each value in the graph below is the percentage of sections (performed in 50 section increments) showing each feature. Blue circles indicate sections where at least part of the pericyte was interacting closely with the EC; purple squares indicate the presence of a ‘peg and socket’- like interaction; magenta triangles indicate sections where a pericyte process is contacting an EC tight junction. Gray shaded region indicates the location of the pericyte cell nucleus. The last value in (c) (at x=19.6) represents the percentage of 40 sections, and the last value in (d) (at x=29.8) represents the percentage of 45 sections. (e) EC ligand-receptor interaction scores in the cortex and ME with values >100 and p-values <0.01. EC ligand interaction scores with mural cell, astrocyte and tanycyte receptors are shown. Candidate, region-specific ligand-receptor interactions investigated below are highlighted in red (cortex-specific) and blue (ME-specific). (f) Co-immunostaining for Bsg (white) and its receptor Integrin α6 (Itga6, red) in cortex and ME, validating elevated expression of ligand (Bsg) and receptor (Itga6) in cortex. Scale bar 5µm. (g) Co-immunostaining for VEGFR2 (red), Emcn (cyan) and (Glut1, green) in cortex and ME. Scale bar 20µm. (h) Immunostaining illustrating complementary spatial distribution of ligand VEGF (white) and receptor VEGFR2 (red) in ME. Non-ME vessels are labeled in green (Glut1). Scale bar 50µm.

### Novel region-specific genes identified in cECs in the ME and cortex

Direct comparison of cortical cECs with ME-derived *Plvap*-expressing cECs revealed 433 differentially expressed genes (Fig. 1i, Extended Data Fig. 4c-d, Table 1). Several novel region-specific genes were validated by immunostaining: Endomucin (Emcn; Fig. 1j) and Endothelial cell specific molecule 1 (Esm1; Extended Data Fig. 2d) are expressed in ME cECs but not in the cortical vasculature, whereas Insulin like growth factor 1 receptor (Igf1r) and Basigin (Bsg) are expressed in the cortical vasculature but are absent from the ME vasculature (Fig. 1k; Extended Data Fig. 2e).

The differences in gene expression observed between the BBB-containing cECs from the cortex and the *Plvap*-expressing cECs from the ME can be classified into six broad categories (Extended Data Fig. 4c-d), including structural components, such as tight junction genes, which are enriched in cortical cECs; transporters, such as *Mfsd2a* and brain capillary marker solute carrier organic anion transporter family member 1C1 (*Slco1c1*), which are enriched in cortical cECs; and transcription factors, such as lymphoid enhancer binding factor 1 (*Lef1*) and catenin beta 1 (*Ctnnb1*), which are enriched in cortical ECs and are associated with Wnt signaling at the BBB^37^. We confirmed lack of Lef1 activity in the ME using TCF/LEF:H2B-GFP reporter mouse^38^ (Extended Data Fig. 4e). These data are also consistent with previous reports of low Wnt activity in the CVOs^39, 40^. All cortex-specific transcripts mentioned above are associated with the specialized function of BBB-containing cECs, including specialized tight junctions and expression of nutrient transporters^41, 42^. Interestingly, ME-associated *Plvap*-expressing cECs show different expression patterns of VEGF receptors than cortical cECs (Extended Data Fig. 4c-d, Fig. 4). These observations are consistent with previous characterizations of the ME as an area of continuous EC proliferation^43^ and the increased permeability of ME blood vessels^27, 44^. *Plvap*-expressing ME cECs also express a number of genes involved in immune regulation, supporting the idea that the ME may be important in neuroimmune interactions^45^. Taken together, the more than 400 molecular differences in cECs in the ME and cortex indicate that the differences in BBB permeability observed in these regions are at least in part due to differences in cECs.

### Distinct astrocyte subtypes with unique interactions with blood vessels found between the ME and cortex

We next compared astrocytes between the ME and cortex. To visualize astrocytes, we used the Ai14 LSL-tdTomato reporter driven by inducible GLAST-CreER (Fig. 2a, Extended Data Fig. 5a-b). Transcript solute carrier family 1 member 3 (*Slc1a3*), which encodes GLAST, is expressed in astrocytes (Extended Data Fig. 5f). Three-dimensional modeling of astrocytes after U.Clear shows that cortical astrocytes were large, stellate cells with their cell bodies situated apart from the vasculature and extending numerous processes that wrapped around blood vessels (Fig. 2a, Extended Data Fig. 5b, Supplementary Video 3). In contrast, the ME contains two different GLAST+ astrocyte subtypes (Fig. 2a, Extended Data Fig. 5b, Supplementary Video 3). One was nestled between ME blood vessels (‘vessel-associated’) with few, very short processes while the other had a cell body near the ventricle (‘ventricle-associated’) with long processes that generally extended towards the ME vasculature. Although the GLAST reporter can also occasionally label tanycytes, we are confident these cells are astrocytes: tanycytes express *Slc1a3* at a lower level than astrocytes and can be clearly distinguished from astrocytes by their morphology and by immunostaining with the tanycyte marker Vimentin (Fig. 2a, Extended Data Fig. 5d-f, Supplementary Video 3).

Accompanying these striking differences in astrocyte cell morphology and distribution, subclustering analysis of astrocytes identified distinct subtypes in each region (Fig. 2b): two cortical subclusters (cortex 1 and cortex 2), two ME subclusters (ME 1 and ME 2) and a subcluster expressing a high level of glial fibrillary acidic protein (*Gfap*) that was predominantly from the ME (‘*Gfap*-high’; Fig. 2b). A number of region-specific transcripts were identified for each astrocyte subcluster (Fig. 2c, Table 1), consistent with previous scRNAseq reports of regional astrocyte heterogeneity in the mouse brain^46, 47^.

To determine which of these newly identified ME astrocyte subtypes represented the ventricle-associated and vessel-associated astrocytes, we utilized our scRNAseq data showing that ME 1 astrocytes express aldehyde dehydrogenase 1 family member L1 (*Aldh1l1*, Fig. 2c). Using *Aldh1l1*-EGFP reporter mice, we find that *Aldh1l1*-expressing astrocytes in the ME are a ventricle-associated astrocyte subtype, which extend few processes toward the ME vasculature (Fig. 2d). In contrast, using *Gfap*-EGFP mice^48^, we found that astrocytes with strong expression of *Gfap* are located in the area between the ventricle and ME blood vessels, extending numerous processes that were in touch with the ME vasculature (Fig. 2e, 3f, Supplementary Video 3), indicating that *Gfap*-high ME astrocytes are also ventricle-associated astrocytes. As both ME 1 and *Gfap-*high astrocyte subtypes are ventricle-associated astrocytes, it is plausible that the ME 2 subcluster represents vessel-associated ME astrocytes. Finally, we also found that *Slco1c1* is highly expressed in cortical astrocytes (Fig. 2c, g) and cortical cECs (Fig. 1i) but is absent in ME astrocytes (Fig. 2c, g) and ME cECs (Fig. 1i) using *Slco1c1-*CreER; Ai14 reporter mice^49^ (see also Extended Data Fig. 5c).

In summary, we observe three ME astrocyte subtypes and two cortical astrocyte subtypes that contact nearby blood vessels in distinct ways. While one ME subtype (likely ME 2) is directly associated with ME blood vessels, the other two subtypes are further away and extend long processes toward the vasculature (ME 1 and Gfap-high). While we observe two cortical subclusters by scRNAseq, morphological characterization reveals that both of these subtypes appear to contact the cortical vasculature through their endfeet. These findings suggest that local EC-astrocyte communication may play an important role in BBB functional specificity. Indeed, we found that most of the top 100 differentially expressed genes between ME and cortical astrocytes are predicted to be secreted or associated with the cell membrane (75% and 69%, respectively; Table 1). This suggests that molecular differences in astrocyte subtypes may be related to intercellular signaling (elaborated in Fig. 4), further supporting the idea that astrocyte-to-EC communication may be important for regional regulation of BBB permeability.

### Pericytes associated with cortex and ME blood vessels show distinct morphological and anatomical features

Although brain pericytes have generally been viewed as a homogenous population, we found several significant differences between ME and cortex pericytes. First, we performed a molecular comparison of pericytes in these regions. We analyzed our scRNAseq data from pericytes together with smooth muscle cells and perivascular fibroblasts because of their similar gene expression profiles^13^ (Fig. 4a). Consistent with previous studies, we found very few significant molecular differences between ME and cortex pericytes.

However, we observed striking morphological differences in pericytes in these two regions as visualized by three-dimensional imaging and reconstruction (Fig. 3b-c, Extended Data Fig. 6f-g, Supplementary Video 4). Sparse labeling of pericytes in platelet-derived growth factor receptor beta (*Pdgfrb*)-CreERT2; Ai14 tdTomato reporter mice revealed that cortical capillary-associated pericytes showed a characteristic ‘bump on a log’ morphology – a prominent nucleus with long processes extending along the length of the vessels (see also Fig. 4) – while ME capillary-associated pericytes have a more irregular shape, a less defined nucleus and processes of varying lengths. Moreover, some ME pericytes contacted more than one capillary whereas cortical pericytes generally interacted closely with a single vessel (Fig 4b-c, Extended Data Fig 6f-g).

We also found different levels of pericyte coverage in these two regions. We used platelet derived growth factor receptor alpha (*Pdgfra*)*-*H2B-GFP; *Pdgfrb*-CreERT2; Ai14 tdTomato reporter mice to quantify the relative number of tdTomato+, GFP-pericyte cells per EC in the ME and cortex (tdTomato+, GFP+ cells are fibroblasts). This analysis revealed approximately half as many Pdgfr*β*-expressing, Pdgfr*α*-negative cells per EC in the ME than in the cortex (Fig. 3d). Previous work indicates that the relative pericyte to EC ratio influences blood vessel permeability^50^, suggesting the lower level of pericyte coverage in the ME may result in increased EC permeability in this region when compared to the cortex. Given that the brain has been reported to have the highest pericyte to EC ratio in the body^50^, it is interesting that the vasculature of the ME, a highly permeable brain region, has locally reduced pericyte coverage when compared to the cortical vasculature.

Finally, we also found that mural and fibroblast cell subtypes were distributed differently in each region in our dataset, despite similar overall cell numbers (Fig. 3e-f, Extended Data Fig. 6e). In the cortex, pericytes were the most prevalent mural and fibroblast cell type sequenced: 98% were mural cells (75% pericytes, 23% SMCs) and 2% were fibroblasts. In the ME, fibroblasts comprised the majority of cells sequenced: 43% were mural cells (20% pericytes, 23% SMCs) and 57% were fibroblasts. Unbiased subclustering analysis of all mural cells revealed pericyte, fibroblast and two smooth muscle cell (SMC) subtypes, which include one SMC subtype with a transcriptional signature of activation (SMCs 2; Fig. 3f). While an ME-specific fibroblast subtype was identified, characterized by expression of transcripts such as decorin (*Dcn*) and collagen type I alpha 1 chain (*Col1a1*; Fig. 3f, Extended Data Fig. 6a-b), region-specific subtypes largely did not emerge.

Taken together, pericytes in the cortex and ME show clear differences in morphology and distribution pattern along capillary vessels, seemingly in the absence of significant gene expression differences. Although it is known that brain pericytes are different from pericytes in the periphery both from developmental origins and gene expression patterns^13, 51, 52^, our data indicate that pericytes are heterogeneous across brain regions. These findings suggest that like astrocytes, pericytes likely contribute to regional differences in barrier properties between the cortex and ME both through direct physical interactions likely also by intercellular communication (elaborated in Fig. 4).

### Serial TEM and U.Clear reveal differences in organization of the vascular and perivascular cellular environment in the ME and cortex

Our three-dimensional fluorescent imaging analyses of ECs, pericytes and astrocytes (Fig. 1, 2, 3) indicate that vascular and perivascular cells are quite different in the cortex and ME. To better understand how these differences translate to functional differences in BBB permeability, we used three-dimensional, whole region imaging and TEM to investigate how these cells interact in each region. Using U.Clear, we co-immunostained for ECs (CD31), pericytes (Pdgfr*β*) and astrocyte endfeet (aquaporin 4 (Aqp4)). Cortical ECs, pericytes and astrocyte endfeet interact closely (Extended Data Fig. 7a). In contrast, ME pericytes are intermingled with ECs in the perivascular space (Extended Data Fig. 7a). Aqp4 is found mainly in ventricle-associated ME astrocytes; however, unlike in the cortex, Aqp4 is not polarized at endfeet but rather found throughout processes that extend toward the ME vasculature (Extended Data Fig. 7b). Thus, there are clear regional differences in how ECs, pericytes and astrocytes physically interact *in situ*.

To gain a better understanding of these interactions at the subcellular level, we performed conventional and serial TEM. Under conventional TEM, cortical capillaries, in addition to their specialized tight junctions and low levels of vesicles, showed close interaction of ECs with pericytes and astrocyte endfeet (Fig. 4a, Extended Data Fig. 8a). As in the cortex, ME cECs shared a basement membrane with pericytes (Fig. 4b, Extended Data Fig. 8b). However, ME cECs were fenestrated with tight junction complexes that appeared shorter than those in the cortex (Fig. 4b, Extended Data Fig. 8b). Strikingly, ME cECs interacted with many fibroblast-like cells in a large perivascular space filled with extracellular matrix protein (Fig. 4b, Extended Data Fig. 8b-c). This is in contrast to the cortical fibroblast-like cells recently described near aECs^13^. The ME parenchyma (highlighted in yellow) abuts ME blood vessels on the dorsal side without reaching into the perivascular space (Fig. 4b, Extended Data Fig. 8b) and contains ME astrocyte and tanycyte processes, which we observed contacting the ME vasculature using three-dimensional fluorescent imaging (Fig. 2, Extended Data Fig. 5).

In light of these distinct, region-specific interactions between pericytes and cECs, we further examined the interaction of cortical ECs and pericytes using serial TEM (Fig. 4c, d, Extended Data Fig. 8d). We reconstructed pericytes and ECs in two capillary blood vessels from a published dataset of the mouse visual cortex^53^. Vessel 1 (Fig. 4c) was reconstructed across 31.8 μm, while vessel 2 (Fig. 4d) was reconstructed across 19.6 μm. These three-dimensional representations reveal how pericyte processes extend from the pericyte cell body along the length of the blood vessels, wrapping around them and sending out smaller processes (Fig. 4c, d, Supplementary Video 5). We quantified the presence of three interaction features along the length of the two blood vessels (Fig. 4c, d). First, we determined the percentage of sections where at least one close pericyte-cEC interaction, defined as where the ECM was not visible between a pericyte process and the EC, was present. In both vessels, a close pericyte-cEC interaction was observed in the majority of sections (in 72% of sections in vessel 1 and 88% of sections in vessel 2; Fig. 4c, d). Next, we analyzed ‘peg and socket’ interactions, where membrane invaginations between ECs and pericytes are prominent. These interactions were rare in vessel 1 (13.8% of sections) but appeared more frequently in vessel 2, particularly in the latter half of the vessel (46% of sections; Fig. 4c, d). Finally, we quantified sections where pericytes contacted EC tight junction clefts, finding this interaction more frequently in vessel 2 (64% of sections) than in vessel 1 (25% of sections; Fig. 4c, d). Taken together, these findings reveal in detail how pericytes and ECs interact along the length of a cortical capillary. The close association of pericytes with ECs and the prevalence of pericyte interactions with EC tight junction clefts, a proposed site of BBB signaling, suggest that these interactions are important for EC-pericyte signaling. In contrast, ‘peg and socket’ interactions were observed less frequently in our data and seemed to be concentrated in smaller domains.

Thus, this new view of ECs and perivascular cells both with EM resolution and using U.Clear has revealed distinct patterns of how perivascular cells interact with ECs within their local environments. These direct, physical interactions and the close proximity of ECs with perivascular cells, particularly in the cortex, further support the possibility that direct signaling from perivascular cells to ECs is essential for BBB maintenance and provide a window into how ligands and receptors may naturally interact to promote intercellular signaling.

### Bioinformatic method identifies novel ligand-receptor pairs between ECs and perivascular cells

To identify unique ligand-receptor interactions between cECs and perivascular cells in the cortex and ME, we performed bioinformatic analyses of our data using a method developed by Kumar and colleagues^54^ (Extended Data Fig. 7c). To tailor our analysis to identifying key interactions, we supplemented their database of experimentally validated ligand-receptor interactions with predicted interactions of our differentially expressed genes in ECs, mural cells and astrocytes (see Methods). This method identified known ligand-receptor pairs between interacting cell types, such as platelet derived growth factor subunit B (Pdgfb)-Pdgfr*β*, an interaction between ECs and pericytes (Fig. 4e). We further identified 26 and 30 unique ligand-receptor pairs between ECs and mural cells in the cortex and ME, respectively (Fig. 4e, Table 1). Additionally, we identified 26 and 25 unique ligand-receptor pairs between ECs and astrocytes in the cortex and ME, respectively (Fig. 4e, Table 1). We confirmed expression of one candidate cortex-specific EC-astrocyte ligand-receptor pair, *Bsg* and integrin alpha 6 (*Itga6*). Immunostaining shows *Bsg* expression in cortex ECs but not in ME ECs and robust *Itga6* expression in cortical astrocyte endfeet – in addition to ECs – but decreased perivascular expression in the ME (Fig. 4f). Finally, by extending our analysis to other ME cell types, we confirmed ME-specific expression of Vegfr2 (*Kdr*) in ECs and found expression of its ligand – Vegfa – in ME-specific tanycyte cells (Fig 5g, h, Extended Data Fig. 7d). Thus, this analysis enabled us to evaluate ligand-receptor expression patterns to identify pericyte- and astrocyte-derived factors that may act upon ECs to maintain BBB integrity. This method both permits better understanding of EC signaling and provides a molecular handle to enable future strategic investigation of how intercellular interactions regulate regional BBB permeability in ECs, which to date is largely unknown.

## DISCUSSION

Here we performed an unbiased comparison of two brain regions with marked differences in BBB permeability and found key molecular and cellular differences, particularly in vascular and perivascular cells. Moreover, by using scRNA-seq, three-dimensional imaging and TEM together, our work examines how regional cell composition and arrangement and intercellular signaling capacity contribute to regional BBB specialization.

scRNAseq of Plvap-expressing cECs from the ME represents the first time this rare cell population has been profiled as single cells extensively. We found approximately 400 differentially expressed genes between cECs from the cortex and the ME. Moreover, a basic comparison of our data to bulk and single-cell RNA sequencing studies of ECs from the mouse pituitary gland^40^ and the mouse neurohypophysis^55^ reveals overlap but also differences in gene expression (Extended Data Fig. 9), suggesting that there may be regional EC heterogeneity even within the ME and the adjacent pituitary gland.

While our morphological analysis of pericytes showed substantial differences between mural cells in the cortex and ME, these differences did not seem to be due to significant differences in gene expression. This is surprising, given their clear morphological differences and in light of recent studies which demonstrate that the developmental origins of pericytes in these two regions are different (Yamamoto et al. 2017, Girolamo et al., 2021). One possibility is that since that we observe cells from the nearby hypothalamus in our dataset, our ME pericyte population is a mixture of pericytes, some which are associated with the Plvap-expressing capillaries in the ME and some that ensheath neighboring BBB-containing vessels. Due to technical limitations, we were unable to remove this contamination. Therefore, whether the lack of pericyte transcriptional heterogeneity we observe is true biology or the result of undersampling the ME pericyte population remains to be seen. In the future, when markers are available to selectively label the pericytes ensheathing Plvap-positive ECs, a clean comparison between these populations can be performed to more carefully address this question.

Three-dimensional imaging of entire regions using U.Clear and visualization of pericyte-EC interactions with serial TEM reconstruction both contextualized our scRNA-seq results and enabled us to view the BBB in a new way, providing us with a deeper understanding of the morphology and arrangement of cells in their natural brain microenvironments in regions with distinct BBB properties. Imaging with U.Clear of the entire local environment with molecular identity allowed us to uncover that while both of these regions contain ECs, astrocytes and pericytes, the physical interaction of these perivascular cells with the vasculature was quite different in each region. Serial TEM reconstruction and analysis of brain endothelial cell and pericyte interactions allowed us to observe several previously unknown features over the length of BBB-containing cortical blood vessels (∼20-30 µm) that conventional EM (∼40-80 nm sections) failed to observe. We found that ECs and pericytes interact closely in the cortex, often through ‘peg-and-socket’-like interactions and at the EC tight junction cleft. Together, these findings highlight how the local cellular environment may influence regional BBB properties and demonstrates the importance of performing both molecular and morphological characterizations to understand BBB heterogeneity within the brain.

Finally, the platform we developed here can be readily applied to examine other brain regions. Comprehensive characterization of other brain regions with noted differences in BBB properties will further clarify how regional differences in cellular organization and intercellular signaling affects EC properties. Most CNS diseases affect specific brain regions. In order to make treatment of these diseases as region-specific as possible, it is critical to identify both regional differences in the BBB and key, region-specific BBB regulators. Thus, understanding how regional BBB heterogeneity is achieved throughout the brain is an important step towards the development of more effective region-specific targeted therapies.

## SUPPLEMENTARY VIDEOS

**Supplementary Video 1. Sulfo-NHS-Biotin leakage in cortex and ME**

Immunostaining for EC marker CD31 (white) and BBB leakage tracer (Sulfo-NHS-Biotin, magenta) showing no leakage of tracer in cortex and tracer leaking out of vessels in ME.

**Supplementary Video 2. Morphology of capillaries in cortex and ME**

High magnification images of capillaries immunostained for CD31 (white) show morphology of cortex and ME capillaries.

**Supplementary Video 3. Morphology of astrocytes in cortex and ME**

Imaris 3D reconstruction of single astrocytes (red) and tanycytes (yellow, in ME) in cortex and ME labeled with Tomato in *Slc1a3*-CreERT2:Ai14 mouse. Co-staining for EC marker CD31 (white). And Imaris 3D reconstruction of single GFAP+ astrocyte (green) in ME labeled with GFP in *GFAP-*EGFP mouse. Co-staining for EC marker CD31 (white).

**Supplementary Video 4. Morphology of pericytes in cortex and ME**

Imaris 3D reconstruction of single pericytes (red) in cortex and ME labeled with Tomato in *Pdgfrb*-CreERT2:Ai14 mouse. Co-staining for EC marker CD31 (white).

**Supplementary Video 5. Serial TEM blood vessel reconstructions**

3D reconstruction of two blood vessel-pericyte interactions from a serial TEM dataset of the visual cortex, highlighting pericytes (blue), an endothelial cell (green) and the blood vessel lumen (red).

## METHODS

### Mice

All mouse experiments followed institutional and US National Institute of Health (NIH) guidelines and were approved by the Harvard University Institutional Animal Care and Use Committee (IACUC). Mice were maintained on a 12 hour light/12 hour dark cycle. All mice used for analysis were 8 to 14 weeks old unless stated otherwise. Both male and female mice were used in all experiments unless otherwise indicated. The following mouse strains were used: wild type (C57BL/6N, Charles River Laboratories #027), Ai14 (JAX: 007914)^56^, *Aldh1l1*-EGFP (JAX: 026033)^57^, GFAP-GFP (JAX: 003257)^48^, Glast-CreER (JAX: 012586)^58^, TCF/LEF-GFP (JAX: 032577) ^59^, *Cdh5*-CreERT2^60^, *Slco1c1*-CreERT2^61^, *Pdgfrb*-CreERT2 (JAX: 029684)^62^, *Pdgfra-* H2B-EGFP (JAX: 007669)^63^, and *Mfsd2a*^ko^ (MMRRC strain 032467-UCD)^64^.

To label specific cell types, heterozygous Glast-CreER, *Pdgfrb*-CreERT2, *Slco1c1*-CreERT2 or *Cdh5*-CreERT2 mice, respectively, were crossed with homozygous Ai14 mice to generate CreER-dependent reporter mice. Recombination and labeling of respective cell types was induced by 5 subsequent intraperitoneal injections of tamoxifen (1mg/mouse) into 8- to 12-week old mice, and brains were collected two weeks later. For sparse cell labeling, a single dose of 4OH-tamoxifen (0.4mg/mouse) was injected one week prior to analysis.

### U.Clear tissue clearing and immunohistochemistry

U.Clear tissue clearing is a newly optimized protocol (Farrell R., Wang W. and Wu Z, *in submission*) based on the Adipo-Clear framework^65, 66^. Briefly, mice were deeply anesthetized with ketamine/xylazine (100 mg/kg) and subsequently intracardially perfused with cold 4% PFA in PBS. Brains were dissected and fixed overnight in 4% PFA at 4°C. For Claudin-5 and LEF1 stainings, brains were perfused with cold PBS and drop-fixed in cold 100% Methanol overnight before rehydrating brains in a series of 70% Methanol/PBS, 30% Methanol/PBS and PBS. After washing brains in PBS, brains were trimmed into two pieces. First, a 5%5%5 mm cube of somatosensory cortex and second, a similarly sized cube of hypothalamus containing the median eminence. Resulting brain pieces were delipidized by 4 washes (1h, 2h, 4h, overnight) with SBiP buffer (200µM Na2HPO4, 0.08% sodium dodecyl sulfate, 16% 2-methyl-2-butanol, 8% 2-propanol in H_2_O (pH 7.4)) at room temperature. Next, brains were transferred into B1n buffer (0.1% Triton X-100, 2% glycine, 0.01% 10N sodium hydroxide, 0.008% sodium azide in H_2_O) for blocking under nutation at room temperature. On the next day brains in B1n buffer were moved to 37 °C for 1 hour. For immunolabeling, primary antibodies diluted in PTxwH buffer (0.1% Triton X-100, 0.05% Tween-20, 0.002% heparin (w/v), 0.02% sodium azide in PBS) were added to brains, and brains were kept at 37 °C with gentle rocking for two days. Antibodies were washed off in 4 washes with PTxwH (1h, 2h, 4h, overnight). Then brains were incubated with secondary antibodies diluted in PTxwH at 37 °C with gentle rocking for two days and subsequently washed 4 times with PTxwH (1h, 2h, 4h, overnight). For further delipidization, samples were immersed in SBiP buffer four times (1h, 2h, 4h, overnight). Next, brains were washed twice in 0.5mM Na2HPO4 (1h, 2h), twice in PB buffer (16mM Na2HPO4, 4mM NaH2PO4 in H_2_O) (1h, 2h) and finally twice in PTS solution (75% PB buffer, 25% 2,2’-thiodiethanol) (1h, overnight), then equilibrated with histodenz gradient buffer with refractive index adjusted to 1.53 using thiodiethanol. Samples were stored at -20 °C until acquisition.

A polyclonal antibody to the C-terminus of mouse Mfsd2a was generated by New England Peptide (Gardner, MA) using IACUC approved protocols. Rabbits were immunized with a KLH-conjugated peptide (Ac-CSDTDSTELASIL-OH). Generated antiserum was purified using a peptide affinity column. Antibody specificity was validated in *Mfsd2a*^ko^ mice (Extended Data Fig. 1f). The following primary antibodies were used at the indicated dilutions: Rabbit polyclonal anti-Aquaporin 4, Millipore AB3594; RRID: AB_91530 (1:200); Goat polyclonal anti-Basigin/EMMPRIN, R&D Systems AF772; RRID: AB_355588 (1:50); Goat polyclonal anti-CD31, R&D Systems AF3628; RRID: AB_2161028 (1:50); Mouse monoclonal anti-Claudin-5 AF488, Thermo Fisher 352588; RRID: AB_2532189 (1:100); Rabbit polyclonal anti-Collagen 1, Millipore AB765P; RRID: AB_92259 (1:100); Goat polyclonal anti-Decorin, R&D Systems AF1060; RRID: AB_2090386 (1:50); Rat monoclonal anti-Endomucin, Santa Cruz sc-65495; RRID: AB_2100037 (1:100); Rabbit monoclonal anti-ERG, Abcam ab92513; RRID: AB_2630401 (1:100); Rabbit monoclonal anti-ERG AF488, Abcam ab196374; RRID: AB_2630401 (1:100); Goat polyclonal anti-Esm1/Endocan, R&D Systems AF1999; RRID: AB_2101810 (1:50); Rabbit polyclonal anti-GFAP, Abcam ab7260; RRID: AB_305808 (1:200); Chicken polyclonal anti-GFP, Aves GFP-1020; RRID: AB_10000240 (1:200); Rabbit polyclonal anti-GFP, Thermo Fisher A21311; RRID: AB_221477 (1:150); Rabbit polyclonal anti-Glut1, Millipore 07-1401; RRID: AB_11212210 (1:100); Rat monoclonal anti-Icam2/CD102, BD Biosciences 553326; RRID: AB_394784 (1:100); Goat polyclonal anti-IGF1R1 R&D Systems AF-305; RRID: AB_354457 (1:50); Rat monoclonal anti-Itga6, R&D Systems MAB13501; RRID: AB_2128311 (1:50); Rabbit monoclonal anti-LEF1, Cell Signaling 2230; RRID: AB_823558 (1:100); Rabbit polyclonal anti-Mfsd2a, This study J9590; RRID: NA (1:100); Goat polyclonal anti-PDGFRb, R&D Systems AF1042; RRID: AB_2162633 (1:50); Rat monoclonal anti-Plvap/Meca32, BD Biosciences 553849; RRID: AB_395086 (1:100); Rabbit polyclonal anti-RFP, Rockland 600-401-379; RRID: AB_2209751 (1:150); Goat polyclonal anti-VEGF, R&D Systems AF-493; RRID: AB_354506 (1:50); Rat monoclonal anti-VEGFR2/Flk-,1 BD Biosciences 555307; RRID: AB_395720 (1:100); Chicken polyclonal anti-Vimentin, Millipore AB5733; RRID: AB_11212377 (1:200).

The following secondary antibodies were used at a 1:250 dilution: donkey polyclonal anti-goat AF488, Jackson Immuno Research 705-545-147; RRID: AB_2336933; donkey polyclonal anti-rabbit AF488, Jackson Immuno Research 711-545-152; RRID: AB_2313584; donkey polyclonal anti-rat AF488, Jackson Immuno Research 712-545-153; RRID: AB_2340684; donkey polyclonal anti-chicken AF488, Jackson Immuno Research 703-545-155; RRID: AB_2340375; donkey polyclonal anti-goat Cy3, Jackson Immuno Research 705-165-147; RRID: AB_2307351; donkey polyclonal anti-rabbit Cy3, Jackson Immuno Research 711-165-152; RRID: AB_2307443; donkey polyclonal anti-rat Cy3, Jackson Immuno Research 712-165-153; RRID: AB_2340667; donkey polyclonal anti-chicken Cy3, Jackson Immuno Research 703-165-155; RRID: AB_2340363; donkey polyclonal anti-goat AF647, Jackson Immuno Research 705-605-147; RRID: AB_2340437; donkey polyclonal anti-rabbit AF647, Jackson Immuno Research 711-605-152; RRID: AB_2492288; donkey polyclonal anti-rat AF647, Jackson Immuno Research 712-605-153; RRID: AB_2340694; donkey polyclonal anti-chicken AF647, Jackson Immuno Research 703-605-155; RRID: AB_2340379.

For quantification of capillary thickness, three ∼50µm thick 40x confocal stacks of capillaries in cortex and ME per mouse were taken. Blood vessels were labelled using the CD31 antibody. To measure capillary diameter, the area covered by 3 different capillaries in each image was measured and divided by their respective vessel length. The average of these 3 diameters was used as average capillary length for a given image. Each data point in graph represents the average capillary diameter of one image. Data points from the same mice are depicted in same colors. Values are expressed as mean ± SD. Significance was determined using a nested two-tailed t-test.

Endothelial nuclei per vessel length were quantified based on three ∼50µm thick 40x confocal stacks of capillaries in cortex and ME per mouse. All ERG+ EC nuclei were counted and total capillary length was measured. All analysis was performed blinded. Each data point in the graph represents an individual image. Data points from same mice are depicted in same colors. Values are expressed as mean ± SD. Significance was determined using a nested two-tailed t-test.

For quantification of pericyte coverage, three ∼50µm thick 40x confocal stacks of capillaries in cortex and ME per mouse were taken. Endothelial cell nuclei were labelled with ERG antibody. Pericytes (GFP-, Tomato+) were identified using *Pdgfra*-H2B-GFP; *Pdgfrb*-CreERT2:Ai14 mice. All analysis was performed blinded. Each data point in the graph represents an individual image. Data points from same mice are depicted in same colors. Values are expressed as mean ± SD. Significance was determined using a nested two-tailed t-test.

### Light microscopy and image analysis

Cleared and stained brains were analyzed at high resolution with a Leica TCS SP8 confocal microscope. Z-stacks of images were processed and 3D reconstructed with Imaris software (version 9.3.1, Oxford Instruments). Photoshop CC and Illustrator CC (Adobe) were used for image processing.

### Permeability assays using tracer injections

All tracers used in this study were injected into circulation by retro-orbital injection under short isoflurane anesthesia. Brains were dissected 30 minutes after tracer circulation, except for microperoxidase, which was circulated for 10 minutes. EZ-Link Sulfo-NHS-LC-Biotin was used at 0.2mg/g bodyweight, HRP was used at 0.5mg/g bodyweight and microperoxidase was used at 1mg/g bodyweight. To test leakage in brains using Sulfo-NHS-Biotin, mice were perfused with 4% PFA after circulation.

### Single-cell RNA sequencing and analysis

#### Sample isolation and dissociation

For each experimental replicate, brain regions of interest were isolated from five, 9-week-old male mice, pooled and processed together. Male mice were selected for scRNAseq analysis to limit variations in ME cellular composition and gene expression patterns due to the estrous cycle, as the ME is a neuroendocrine region that is responsive to reproductive hormones^67^. Mice were euthanized at 8 AM to avoid variation due to circadian cycle. Samples were collected on 11 separate days. Brain regions of interest were dissociated into single cells using a protocol adapted from Hrvatin and colleagues (2017). Brains were dissected into cold dissociation medium (1X HBSS without calcium and magnesium, 0.01M HEPES, 9 mM MgCl2, 35 mM D-glucose, pH 7.35), and all subsequent microdissection steps were performed in dissociation medium on ice. From each brain, the median eminence was microdissected using deWecker scissors, then the rest of the brain was sectioned into 1 µm thick sections using a brain sectioning matrix (Ted Pella). Samples of the cortex were obtained using a 1 mm tissue biopsy puncher (Harris Uni-Core). Samples were then dissociated using the papain dissociation system (Worthington) according to the manufacturer’s instructions with the following modifications. Dissociation was performed at 37°C for 45 minutes with gentle agitation in the dissociation medium described plus papain at a concentration of 20 U per milliliter. After gradient centrifugation with the ovomucoid inhibitor, cells were washed in dissociation medium containing 0.04% BSA and resuspended in dissociation medium with 0.04% BSA and 15% Optiprep (Sigma) for inDrops cell encapsulation. Single cell RNA-Seq was provided by the Single Cell Core at Harvard Medical School.

#### inDrops library preparation, sequencing and data processing

One or two libraries of approximately 3,000 cells were collected from each experimental sample. inDrops was performed as described previously^33, 34^ using v3 barcodes, and 44 libraries were generated over 8 separate days to minimize variation due to library preparation as much as possible. Libraries were then pooled and sequenced in 18 runs with the NextSeq 500 using the high output flow cell (Illumina), pooling 3,000 to 12,000 cells per sequencing run. Transcripts were processed using the latest version of the inDrops pipeline (https://github.com/indrops/indrops) and mapped to the mouse transcriptome (ENSEMBL release 85). All steps of the pipeline were run with the default parameters. *Quality control and clustering analysis.* Analysis was performed with R^68^ (version 3.4.1) and RStudio^68^ using the Seurat analysis package^35, 36^ (version 2.3.4). Cells were filtered based on number of genes expressed (between 200 and 4,000) and percentage of mitochondrial genes (less than 20%) to select for high quality, single cells. Across the dataset, the average nUMI is 2,510 and the average number of genes is 1,287 per cell. All cells were combined into a single dataset to permit comparative analysis. The dataset was log-normalized and scaled to 10,000 transcripts per cell. Highly variable genes were determined using the FindVariableGene function using the experiment mean and LogVMR dispersion function and the following parameters to set the minimum and maximum dispersion: x.low.cutoff = 0.0125, x.high.cutoff = 3, y.cutoff = 0.5. Data was scaled to regress out the following variables: number UMIs, percent mitochondrial genes, number of genes, library batch and experimental run date. Next, principal component analysis was performed using the highly variable genes. The data was then clustered using the top 50 principal components with the clustering resolution set to 0.6, resulting in 41 clusters. We obtained 101,347 single-cell transcriptomes that passed initial quality control metrics.

Several region-specific clusters emerged. An ME-specific EC cluster (ECs 2), characterized by expression of *Plvap* (Figs. 1f, g, Extended Data Fig. 2c) is the rare cell population that corresponds to the ME-specific ECs we observed by immunostaining (Fig. 1). As expected, neurons from each region generally clustered separately, and cell types that are unique to the ME such as tanycytes and pars tuberalis cells were derived only from the ME sample (Fig. 1g, Extended Data Fig. 2c). *Cell type assignment and subclustering analyses.* Marker genes for each cluster were determined with the FindAllMarkers function using the default Wilcoxon rank sum test and the following parameters: only.pos = TRUE, min.pct = 0.25, thresh.use = 0.25. To assign general cell types to each cluster, known marker genes were used: *Nrgn, Tubb3, Syt1, Snap25, Camk2b, Thy1* (neurons), *Cldn11, Mog, Mbp, Plp1, Mobp, Opalin, Olig1* (oligodendrocytes); *Olig1, Olig2, Pdgfra, Gpr17, Cspg4* (OPCs); *Aqp4, Aldoc, Slc1a2, Agt, Gja1, Atp1a2, Glul, Vegfa* (astrocytes); *Gpr50, Rax, Crym, Vegfa, Vim, S100a6* (tanycytes), *Cdh5, Flt1, Ptprb, Tek* (ECs); *P2ry12, C1qa, C1qb, Csf1r, Cx3cr1, Itgam, Tmem119, P2ry13* (microglia); *Rgs5, Notch3, Pdgfrb, Cspg4, Vtn, Acta2, Kcnj8, Abcc9, Myh11* (mural cells); *Cd74, Lyve1, Mrc1, Ptprc, H2.Eb1, H2.Ab1, Adgre1* (PVMs), *Apoc3, Chga, Cga, Chgb, Cck, Timeless, Tshb, Cyp2f2* (pars tuberalis cells), *Dcn, Col1a1, Col1a2, Col3a1, Lum, Pdgfra* (fibroblasts); *Ccdc153, Rarres2, Vim, Tmem212, Foxj1* (ependymal cells), and *Ccl5, Ptprc, Trbc2, Cd52, Il2rb* (T cells). General cell types were further analyzed to remove doublets and reveal cell type subclusters. To best resolve subclusters, ECs, astrocytes, neurons, oligodendrocytes and pars tuberalis cells were analyzed individually, while tanycytes and ependymal cells, mural cells and fibroblasts, and microglia, PVMs and T cells were analyzed together. For each subclustering analysis, subclusters expressing marker genes characteristic of multiple cell types were considered doublets and removed, then subclustering was performed again. After subclustering analysis was complete (detailed below), all cells were analyzed together again using the same parameters described above to generate the final dataset containing 65,872 cells and 34 clusters.

#### Classification of cell subtypes

For each subclustering analysis, clustering analysis was performed as for the complete dataset described above. Details about cell subtype assignment for ECs, astrocytes and mural cells and fibroblasts are described below. In addition to vascular and perivascular cell subtypes, we identified 18 subtypes of neurons, including 8 that were ME-specific and 8 that were cortex-specific. Consistent with previous reports^24, 46, 69^, we discerned several subytpes of oligodendrocytes and tanycytes. Finally, we identified several immune cell subtypes^70, 71^, which include macrophages, PVMs and 3 subclusters of microglia.

<u>ECs</u>: Subclustering analysis of EC clusters revealed 6 subclusters. One cluster expressed marker genes characteristic of both ECs and pericytes (described above). The cells in this cluster were considered doublets and were removed from subsequent analyses. The remaining clusters expressed marker genes characteristic of the segments of the vasculature or *Plvap*. The cECs 2 subcluster shows a gene expression signature characteristic of activation, which may be related to activity-induced transcriptional responses in ECs^72^ or activation following tissue disscociation^73^. We also compared gene expression patterns in aECs and vECs between these regions, revealing 13 and 8 differentially expressed genes in aECs and vECs, respectively (Table 1). Consistent with the notion that the BBB resides in cECs, these genes do not encode tight junction components or transcytosis regulators but are likely related to other physiological differences between the ME and cortex. Nevertheless, we were still able to identify a small number of regionally enriched genes in larger blood vessels, further highlighting the ability of our approach to identify transcriptional heterogeneity within brain ECs. <u>Astrocytes</u>: Subclustering analysis of astrocytes revealed 5 subclusters. One astrocyte cluster, predominantly comprised of cells from the ME, expressed high levels of *Gfap*. 2 additional astrocyte subclusters were isolated from the ME, in addition to 2 subclusters from the cortex. <u>Mural cells and fibroblasts</u>: Subclustering analysis of mural cells and fibroblasts revealed 4 subclusters, which correspond to fibroblasts, pericytes and 2 populations of SMCs. Subclustering did not reveal region-specific pericyte cell subclusters, so we compared pericytes from the ME and cortex to find regional differences in gene expression. This analysis identified 5 genes that were differentially expressed in pericytes between the cortex and the ME using a corrected p-value (Table 1). <u>Microglia, PVMs and T cells</u>: Subclustering analysis of immune cells revealed 3 subclusters of microglia, a macrophage subcluster, a PVM subcluster, a T cell subcluster and a B cell subcluster. <u>Tanycytes and ependymal cells</u>: Subclustering analysis of tanycytes and ependymal cells revealed 8 subclusters, which correspond to ependymal cells, *α*2 tanycytes, 2 populations of *α*1 tanycytes, 2 populations of *β*1 tanycytes and 2 populations of *β*2 tanycytes. <u>Oligodendrocytes</u>: Subclustering analysis of oligodendrocyte clusters revealed 10 subclusters, which include OPCs, proliferating oligodendrocytes, newly formed oligodendrocytes, and 7 populations of mature oligodendrocytes. <u>Neurons</u>: Subclustering analysis of neuron clusters revealed 18 subclusters. <u>Pars tuberalis</u>: Subclustering analysis of pars tuberalis cells revealed 3 subclusters.

#### Differential gene expression analysis and comparison to pituitary gland and neurohypophysis ECs

Differentially expressed genes between groups of cells were determined using the FindMarkers function, using the default parameters (Wilcoxon rank sum test). Functional categories of differentially expressed genes were determined using the Uniprot database^74^. The intersection between the top 100 marker genes of the ME-derived, Plvap-expressing cECs (this study; Table 1 – page 3), the top 100 marker genes from the vascular endothelial cells from the scRNAseq dataset of the mouse neurohypophysis^55^, and the top 100 genes expressed in pituitary ECs^40^ is displayed with a Venn diagram. The 21 genes common to all datasets are *Plvap, Igfbp3, Plpp3, Plpp1, Col15a1, Ldb2, Exoc3l2, Cd24a, Meis2, Btnl9, Ltbp1, Cd300lg, Ehd4, Tnfaip2, Ramp3, Slc43a3, Gpihbp1, Cyp26b1, Fabp4, Irx3,* and *Prss23*.

### Transmission electron microscopy

Adult mice were anesthetized with ketamine/xylazine via intraperitoneal injection and then transcardially perfused with cold 5% glutaraldehyde, 4% PFA and 0.1M sodium cacodylate. Brains were dissected and post-fixed by immersion in 5% glutaraldehyde, 4% PFA and 0.1M sodium cacodylate overnight at 4°C. Following fixation, brains were washed overnight in 0.1 M sodium cacodylate.

Brains were washed 3×15 minutes in 0.1M sodium cacodylate, then sectioned into 100 μm free-floating coronal sections with a vibratome. Regions of interest were microdissected, post-fixed in 1% osmium tetroxide and 1.5% potassium ferrocyanide, dehydrated and embedded in epoxy resin. Ultrathin sections of 80 nm were then cut from the block surface, collected on copper grids and counter-stained with Reynold’s lead citrate before examination under a 1200EX electron microscope (JEOL) equipped with a 2k CCD digital camera (AMT).

### Serial transmission electron microscopy reconstruction

Serial transmission electron microscopy data of the visual cortex was generated previously in a 450 μm × 450 μm × 50 μm volume^53^. Three-dimensional rendering of the cortical vasculature consisted of tracing the interaction of ECs and pericytes in FIJI ^75^ using TrakEM2 ^76^. First, the EC was identified by its large central nucleus, thin membrane and proximity to a blood vessel lumen. Next, an interacting pericyte was identified by its perivascular location and its cytoplasmic processes interacting with the EC. Each cell type was manually traced in each section in the dataset, then the images were rendered together to create a three-dimensional reconstruction. Vessel 1 was reconstructed through 794 serially cut 40nm sections, with 66 sections excluded. Vessel 2 was reconstructed in 490 serially cut 40 nm sections, 28 sections excluded. Three-dimensional images were processed and rendered in blender (blender.org).

### Ligand-receptor analysis

To investigate intercellular signaling patterns, an analysis similar to that by Kumar and colleagues^54^ was performed. First, a ligand-receptor database was assembled. To facilitate discovery of novel ligand-receptor pair interactions, the curated reference database the authors used previously was supplemented. The predicted subcellular localization of differentially expressed genes in each candidate interacting cell type was determined using the Uniprot database^74^. For those genes that were known or predicted to be present in the plasma membrane, secreted proteins or components of the ECM, the STRING database was queried to identify candidate interaction partners^77^. Interaction partners with experimental validation were added to our ligand-receptor database. Next, an interaction score was calculated for each ligand-receptor pair for two candidate interacting cell types of interest by multiplying the average expression of the ligand gene in the candidate signaling cell type and the average expression of the receptor gene in the candidate receiving cell type. Finally, the significance of observing a particular interaction score in our dataset was determined by bootstrapping. Briefly, for each iteration of the bootstrapping estimation, the average expression of all genes in each identified cluster was calculated. The dataset was randomized with replacement and interaction scores were calculated for 2,192 random pairs (the size of the supplemented database described above). This iteration was repeated 1000 times to generate a null distribution of interaction score values. From this distribution, we determined that values greater than 30 were statistically significant, as these values were observed with a one-sided p-value less than 0.01 after Bonferroni correction for multiple comparisons (Table 1).

## ACKNOWLEDGEMENTS

The authors would like to thank members of the Gu laboratory for comments on this manuscript; the Single Cell Core at HMS for performing the single cell RNA-seq sample preparation; Elio Raviola for advice and discussion; Sinisa Hrvatin, M. Aurel Nagy and Rory Kirschner for advice and technical assistance; Markus Schwanniger for providing *Slco1c1*-CreER mice; and Ralf Adams for providing *Cdh5*-CreERT2 mice. Imaging, consultation and/or services were in part performed in the Neurobiology Imaging Facility. This facility is supported in part by the HMS/BCH Center for Neuroscience Research as part of an NINDS P30 Core Center grant (NS072030). Electron Microscopy imaging, consultation and services were performed in the HMS Electron Microscopy Facility. S.J.P is a Damon Runyon Fellow supported by the Damon Runyon Cancer Research Foundation (DRG-2289-17). U.H.L. is an EMBO long-term fellow supported by an EMBO long-term fellowship (ALTF 42-2017). R.A.L. is supported by the Howard Hughes Medical Institute Exceptional Research Opportunities Program. The research of C.G. was supported in part by a Faculty Scholar grant from the Howard Hughes Medical Institute. This work was also supported by the Fidelity Biosciences Research Initiative (C.G.), an Allen Distinguished Investigator Award, a Paul G. Allen Frontiers Group advised grant of the Paul G. Allen Family Foundation (C.G.), the National Institute of Neurological Disorders and Stroke (R35NS116820 and DP1NS092473 to C.G.) and the National Cancer Institute (DP1NS092473 to C.G.). Research reported in this publication was also supported by the National Institute On Drug Abuse of the National Institutes of Health under Award Number RF1DA048786 to C.G. The content is solely the responsibility of the authors and does not necessarily represent the official views of the National Institutes of Health.

## AUTHOR CONTRIBUTIONS

S.J.P, U.H.L and C.G. conceived the project, designed the experiments and wrote the manuscript with input from all authors. S.J.P. and U.H.L performed experiments and data analysis. T.F., I.P. and F.N. performed three-dimensional reconstruction of serial TEM data. R.A.L. performed immunostainings. W.-C.A.L. provided technical guidance for serial TEM analysis. Z.W. provided technical guidance and experimental resources for U.Clear experiments.

## COMPETING INTERESTS

The authors declare no competing interests.

## MATERIALS & CORRESPONDENCE

Correspondence and material requests should be addressed to Chenghua Gu.

---

**Extended Data Fig. 1.**
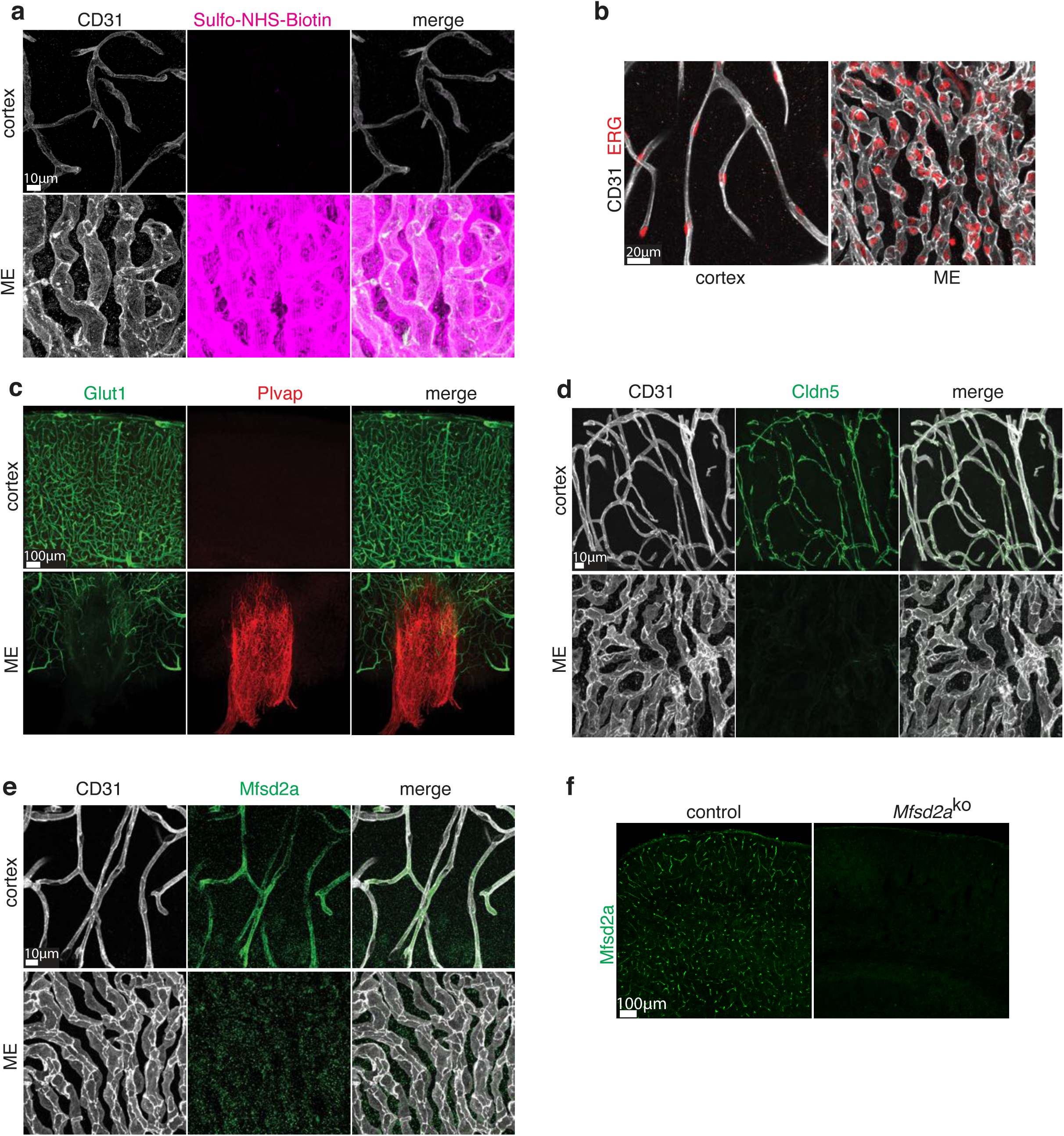
Morphological, molecular and functional differences of the vasculature between the ME and cortex. (a) Tracer leakage assay using Sulfo-NHS-Biotin (magenta) as tracer and immunostaining for blood vessels (CD31, white) in cortex (upper panel) and ME (lower panel). Tracer in circulation has been washed out by perfusion prior to analysis. Scale bar 10µm. (b) Representative images of ERG (red) and CD31 (white) staining in cortex and ME used for quantifications in Figure 1 D. Scale bar 20µm. (c) Co-immunostaining of Glut1 (green) and Plvap (red) in ME and cortex. Scale bar 100µm. (d) Co-immunostaining of CD31 (white) and tight junction protein Cldn5 (green) in cortex and ME. Scale bar 10µm. (e) Co-Immunostaining of CD31 (white) and Mfsd2a (green) in cortex and ME. Scale bar 10µm. (f) Validation of specificity of newly generated polyclonal antibody against Mfsd2a (green) by immunostaining of cortex from wild type and *Mfsd2a*^ko^ mice. Scale bar 100µm.

**Extended Data Fig. 2.**
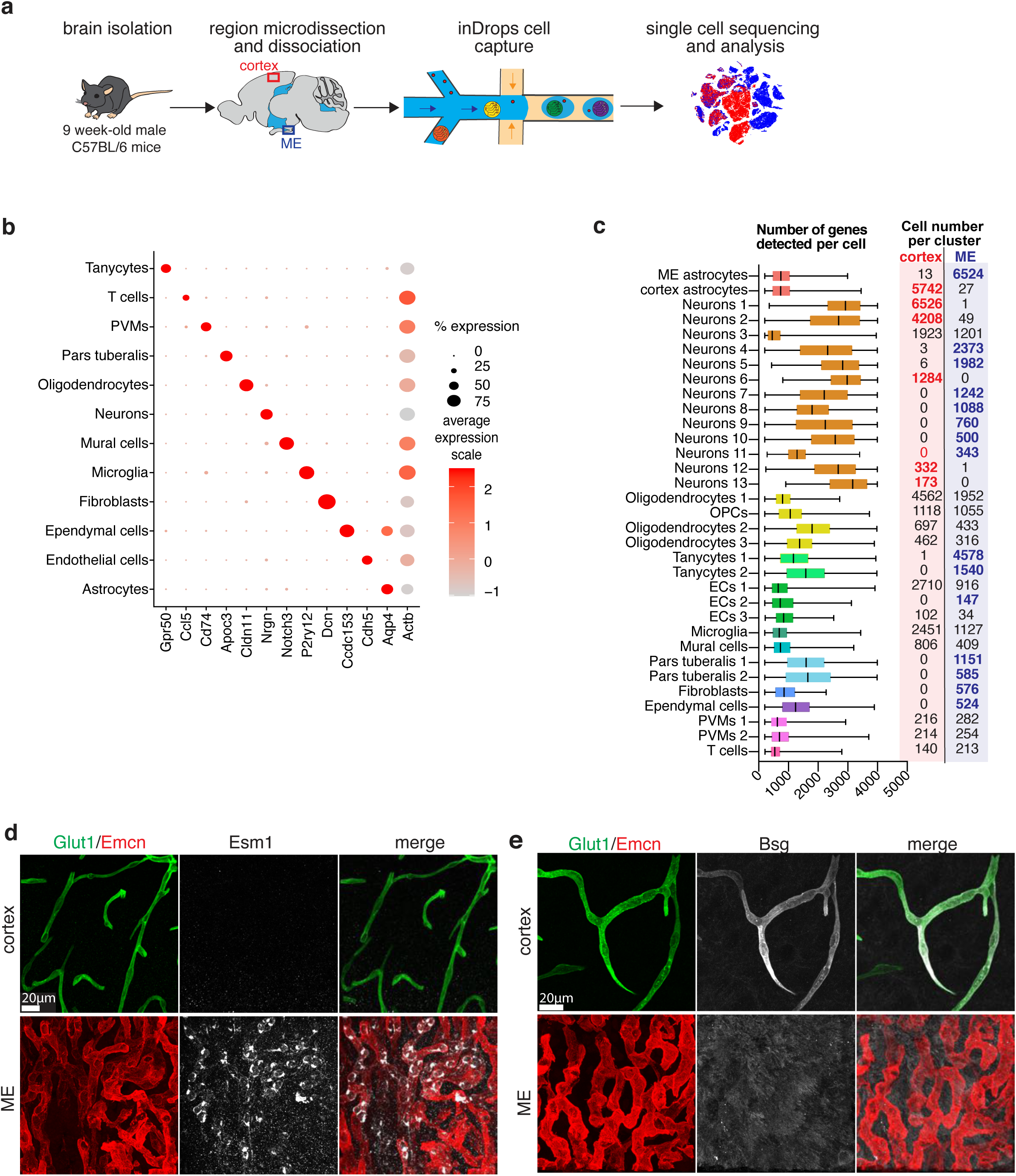
Single cell RNA sequencing of median eminence and a size-matched region of somatosensory cortex reveals unique cell types in each brain region. (a) Overview of the scRNAseq experimental workflow. (b) Dot plot of average expression of cell type specific transcripts used to annotate cluster cell types in Fig. 1f. (c) Box and whisker plot depicting the number of genes detected per cell in all identified clusters in (a). Box indicates median, first and third quartiles, whiskers indicate the minimum and maximum values in each distribution. The cell number in each cluster per sample region is indicated at the right of each plot. (d) Co-immunostaining of Esm1 (white), Emcn (red) and Glut1 (green) in cortex and ME, validating Esm1 is ME cEC-enriched. Scale bar 20µm. (e) Co-immunostaining of Bsg (white), Glut1 (green), and Emcn (red) in cortex and ME, validating that Igfr1 is cortex cEC-enriched. Scale bar 20µm.

**Extended Data Fig. 3.**
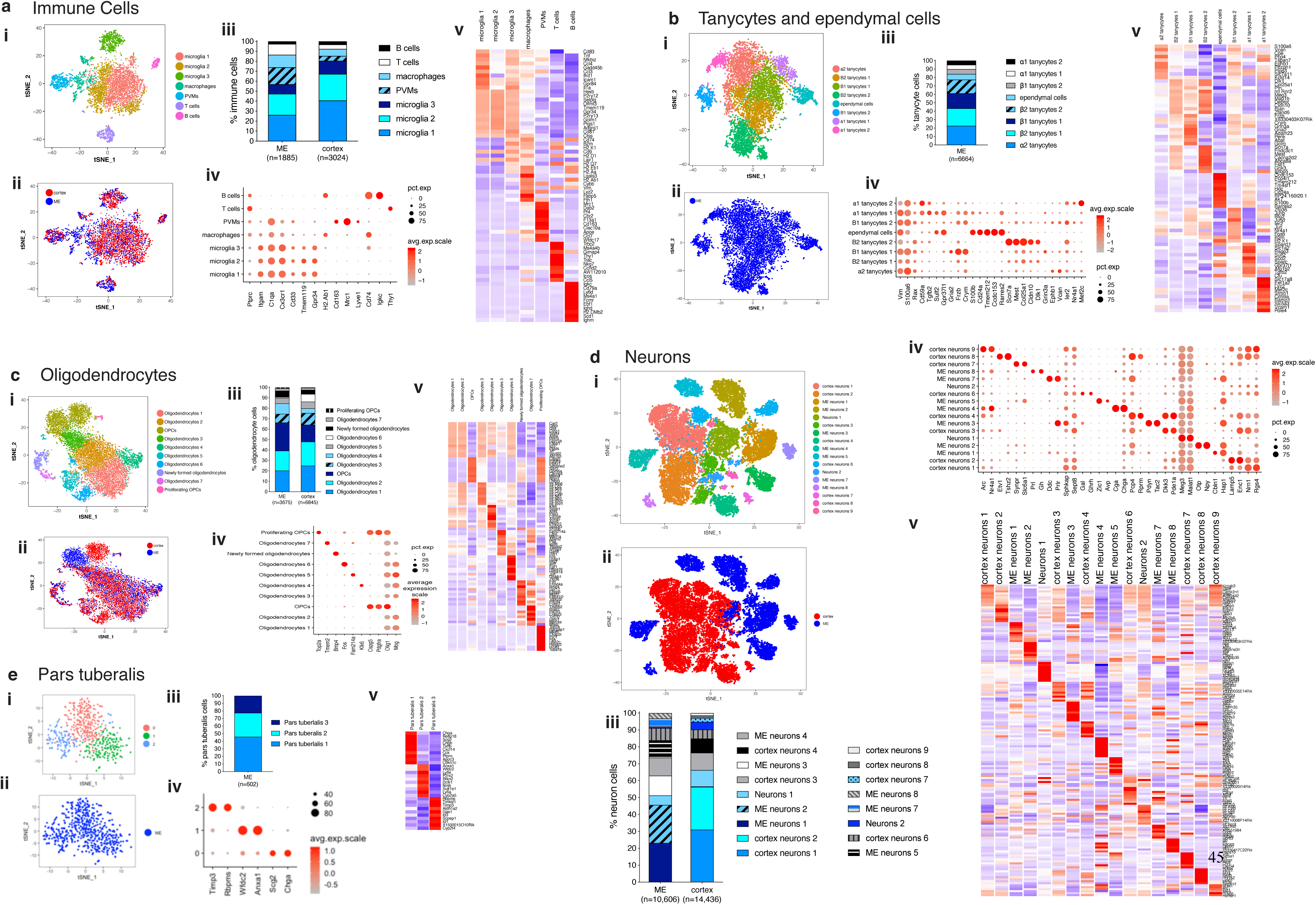
Single cell RNA sequencing of median eminence and a size-matched region of somatosensory cortex reveals unique cell types in each brain region. Subclustering analyses of (a) immune cell, (b) tanycyte and ependymal cell, (c) oligodendrocyte, (d) neuron and (e) pars tuberalis cell types. In each panel there is (i) a tSNE plot of cell type subclusters, (ii) a tSNE plot of cell type subclusters by sample region, (iii) a bar graph showing the distribution of each subcluster within each sample region, (iv) a dot plot displaying marker genes for each subcluster, and (v) a heatmap illustrating the average expression of the top 10 genes in each subcluster. Tanycytes and ependymal cells (b) and pars tuberalis cells (e) were isolated only from the ME sample.

**Extended Data Fig. 4.**
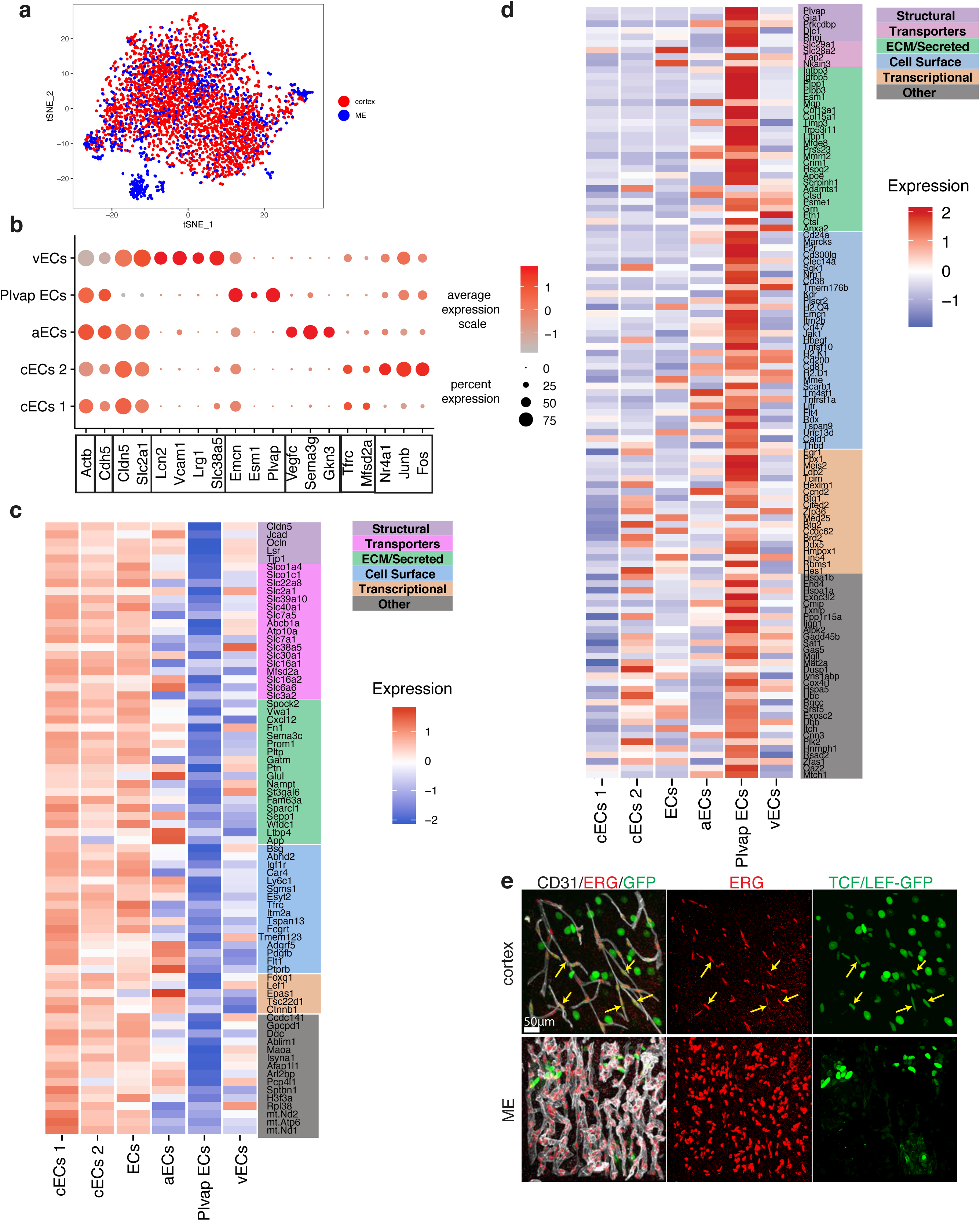
Novel region-specific genes identified in cECs in the ME and cortex. (a) tSNE projection in Fig. 1h colored by sample region. (b) Dot plot of average expression of vascular zonation markers and region-specific transcripts used to annotate cluster cell types in Figure 1h. Genes are color-coded based on EC subtype. *Actb* and *Cdh5* are expressed in all populations, and *Cldn5* and *Slc2a1* are cortex-specific transcripts, as illustrated in Figure 1. (c) Heatmap illustrating the increased relative expression of cortex-derived cEC 1 genes when compared to ME-derived Plvap-expressing ECs. Enriched genes are classified and color-coded based on the type of molecule they encode. (d) Heatmap illustrating the increased relative expression of ME-derived Plvap-expressing EC genes when compared to cortex-derived cECs 1. Enriched genes are classified and color-coded based on the type of molecule they encode. (e) Co-immunostaining for CD31 (white), endothelial nuclei marker ERG (red), and GFP in cortex and ME of TCF/LEF-GFP Wnt-signaling reporter mice. GFP expression (green) indicates activation of the Wnt-signaling pathway. Arrows indicate ERG and GFP double positive nuclei. Scale bar 50µm.

**Extended Data Fig. 5.**
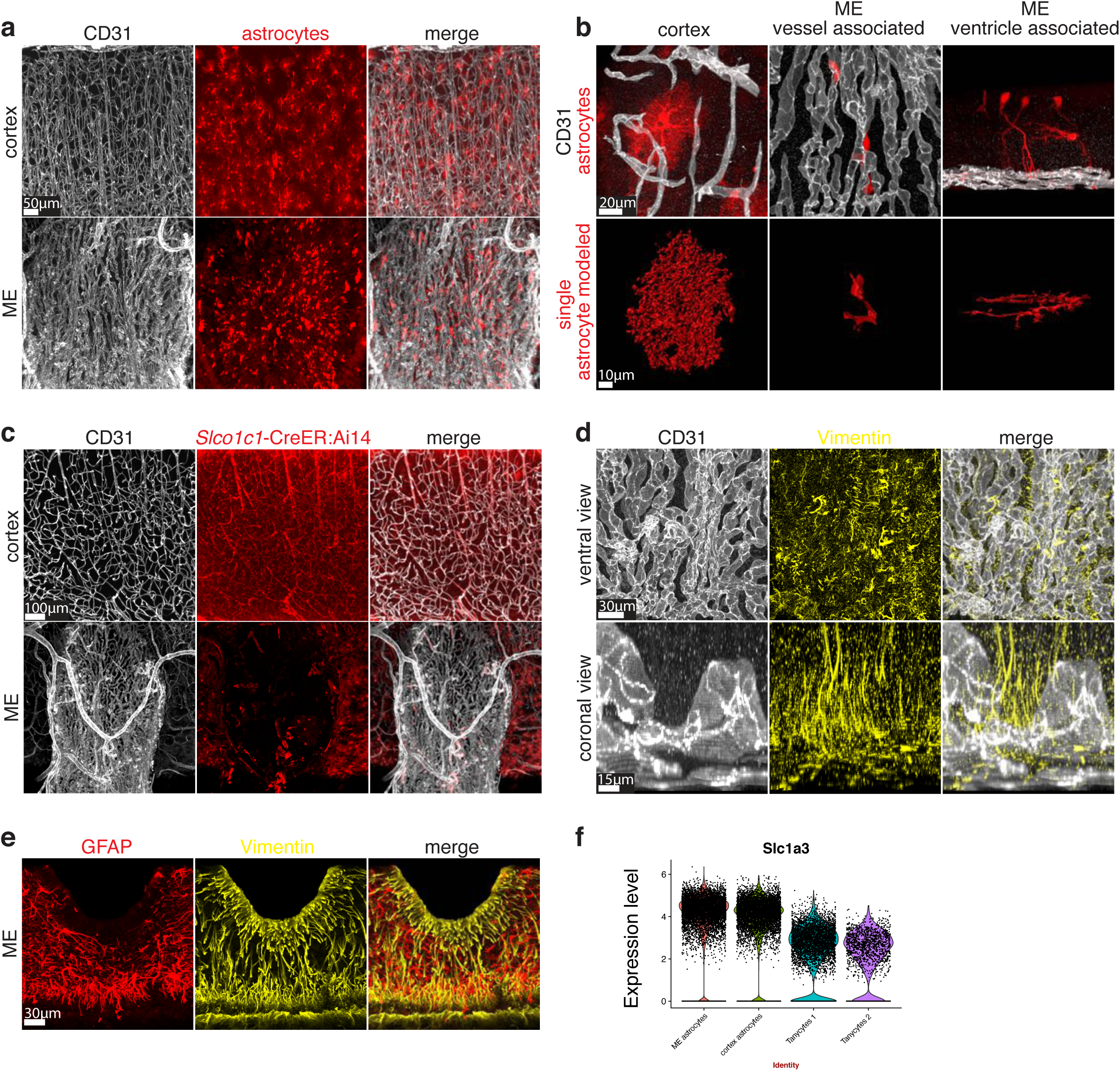
Astrocyte subtypes and their interactions with blood vessels are distinct between the ME and cortex. (a) Fluorescent labeling of astrocyte populations (Tomato, red) in cortex and ME using Glast-CreER:Ai14 mice with a high dose of tamoxifen. Co-staining for blood vessels (CD31, white). Scale bar 50µm. (b) Fluorescent labeling of astrocyte populations in the cortex and ME using Glast-CreER:Ai14 mice after a low dose of tamoxifen to achieve sparse cell labeling. Upper row: immunostaining for Tomato-positive astrocytes (red) and blood vessels (CD31, white). Lower row displays 3D reconstructions of astrocytes (red). Scale bar 20µm (upper) and 10 µm (lower). (c) Fluorescent labeling of cortex blood vessels and astrocyte populations (Tomato, red) in cortex and ME using *Slco1c1*-CreER:Ai14 mice. Co-staining for blood vessels (CD31, white). Scale bar 100µm. (d) Co-immunostaining for tanycyte marker Vimentin (yellow) and pan EC marker CD31 (white) in ME. Ventral and coronal view of ME shown, note tanycyte protrusions in touch with ME vessels. Scale bar 30µm (upper panel) and 15um (lower panel). (e) Co-immunostaining for GFAP-enriched astrocyte population (GFAP, red) and tanycyte marker Vimentin (yellow). Scale bar 30µm. (f) Violin plot showing expression of *Slc1a3,* which encodes GLAST, in astrocyte and tanycyte populations.

**Extended Data Fig. 6.**
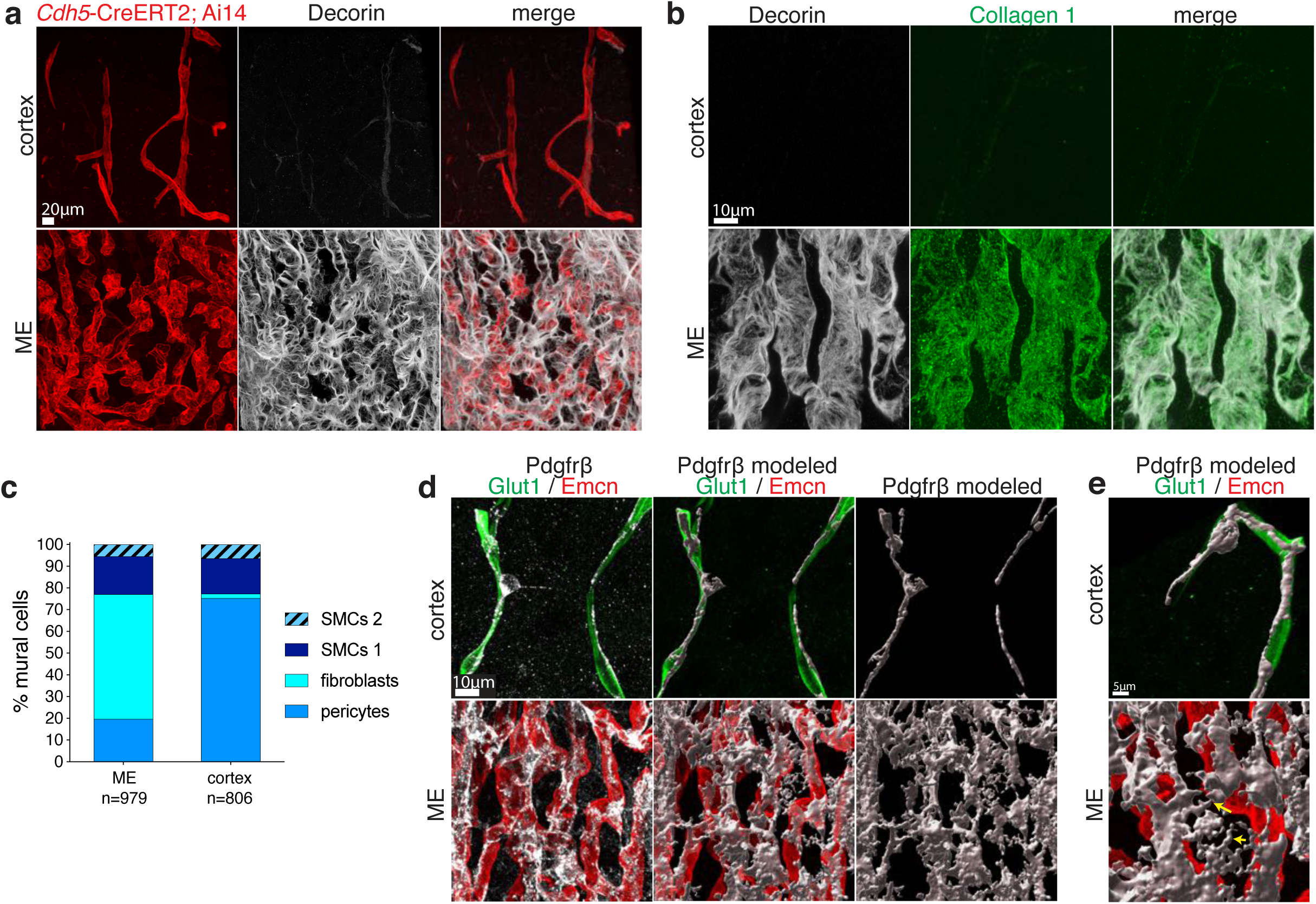
Pericytes associated with cortex and ME blood vessels show distinct molecular, morphological and anatomical features. (a) Co-immunostaining for fibroblast enriched protein Decorin (white) and Tomato in cortex and ME. Blood vessels labelled with Tomato using *Cdh5*-CreERT2:Ai14 mice. Scale bar 20µm. (b) Co-immunostaining for fibroblasts using Decorin (white) and Collagen 1 (green) in cortex and ME. Scale bar 10µm. (c) Bar graph showing the proportion of cells of each cell subtype in the mural cell subclustering analysis in each sample region. (d) Immunostaining for Pdgfrβ (white, left panel) and Imaris 3D reconstruction of pericytes (white, middle and right panel) in cortex and ME. Co-staining for Glut1 (green) and Emcn (red) to mark capillaries. Scale bar 10µm. (e) High magnification images of reconstructed pericytes (Pdgfrβ, white) in contact with capillaries (Glut1, green and Emcn, red) in cortex and ME. Arrows point at ME pericyte protrusions not in contact with capillaries. Scale bar 5µm.

**Extended Data Fig. 7.**
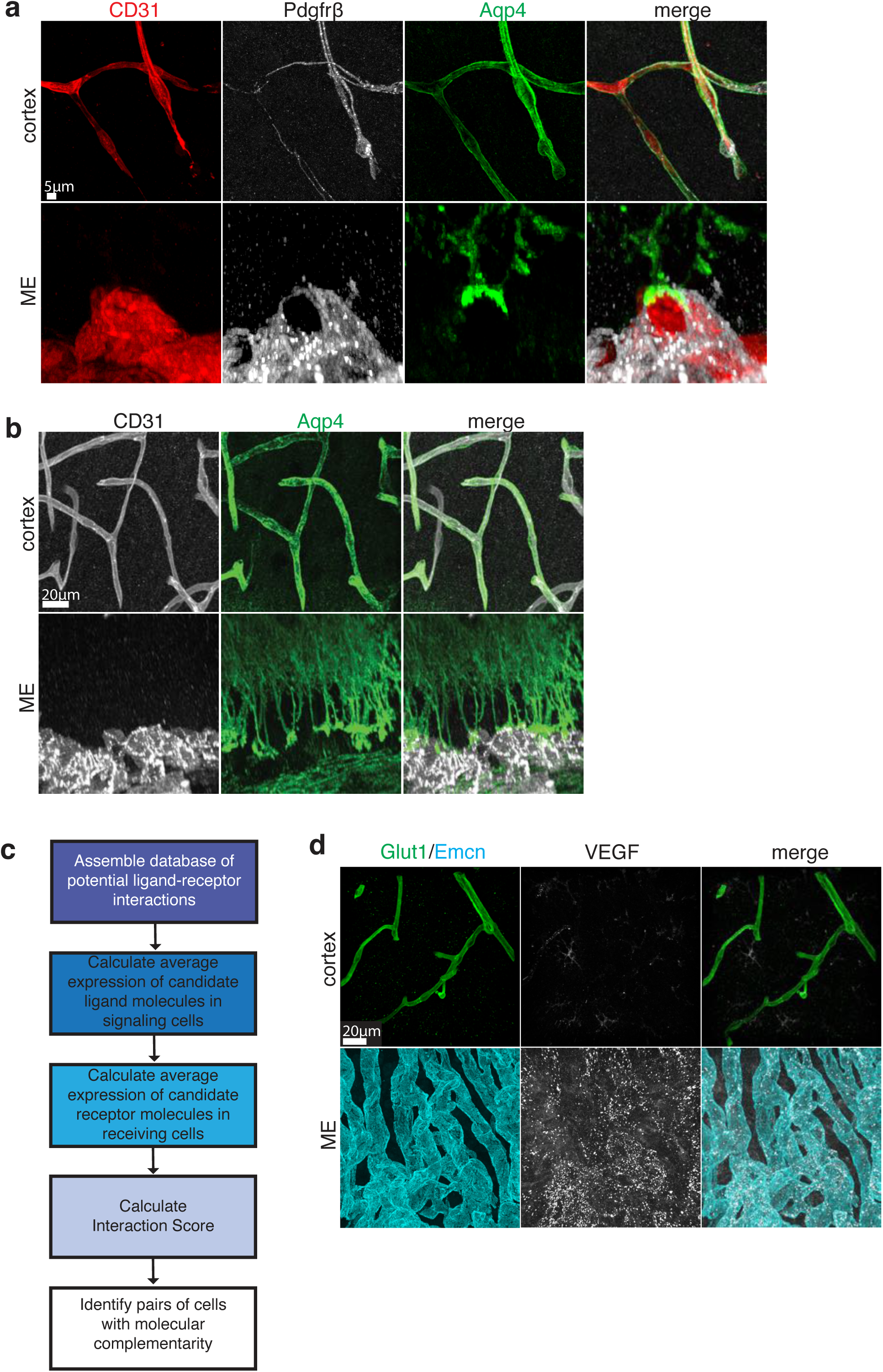
Serial TEM and U.Clear reveal differences in organization of the vascular and perivascular cellular environment in the ME and cortex. (a) Immunostaining for CD31 (red), pan-pericyte marker Pdgfrβ (white) and astrocyte endfoot marker Aqp4 (green) in cortex (left) and ME (right). Scale bar 5µm. (b) Co-immunostaining for astrocyte endfoot marker Aqp4 (green) and CD31 (white) in cortex and ME. Scale bar 20µm. (c) Overview of ligand-receptor analysis methodology and workflow. (d) Co-immunostaining for VEGF (white), Emcn (cyan) and (Glut1, green) in cortex and ME. Scale bar 20µm.

**Extended Data Fig. 8.**
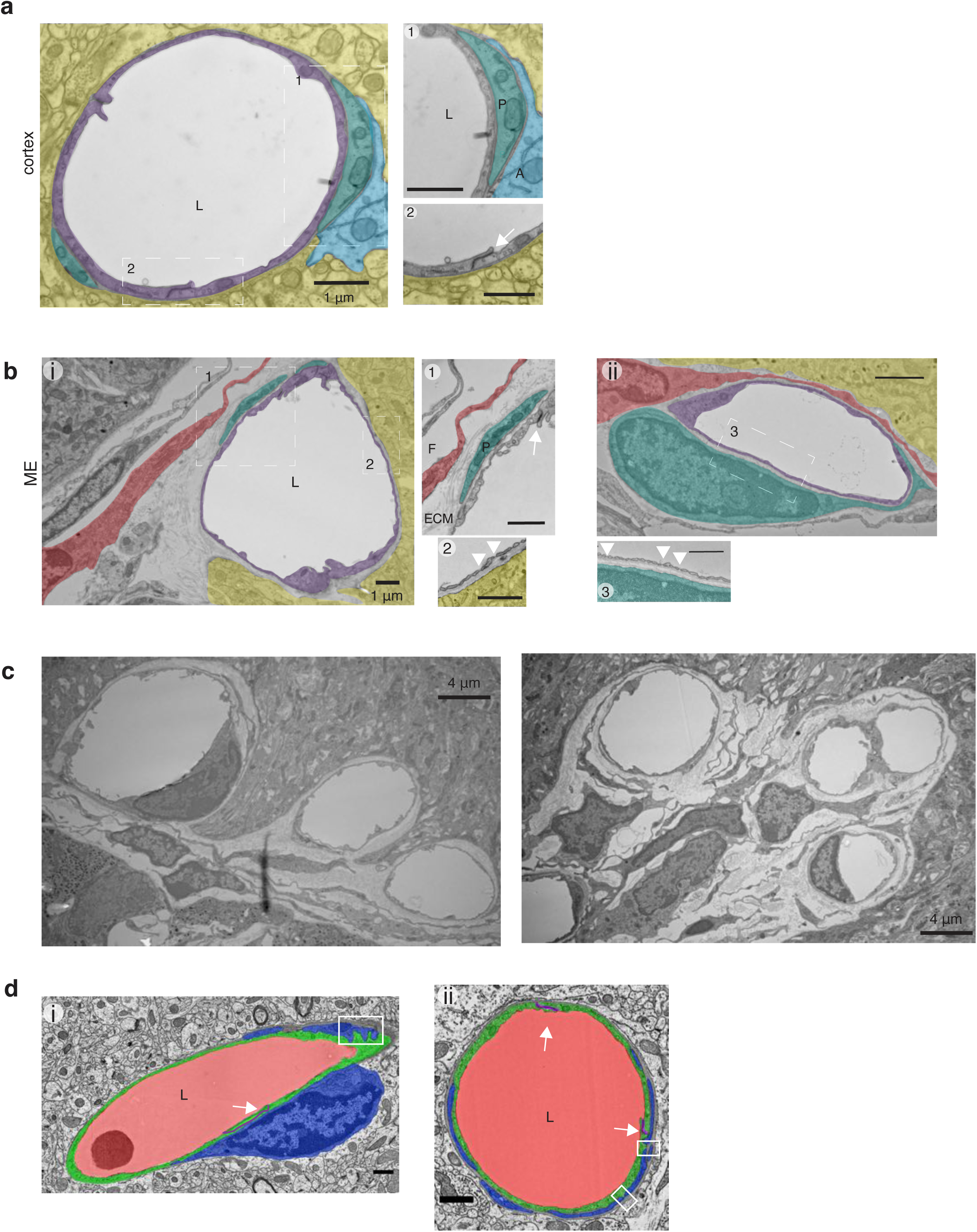
TEM reveals differences in organization of the vasculature and cellular environment in the ME and cortex. (a) TEM images of a cortical capillary. Pseudocolors highlight different cells: cEC (purple), pericyte (teal), astrocyte endfoot (cyan), lumen (L, white) and neuropil (yellow). Insets show cEC tight junctions (white arrows), pericyte cells (P, teal) and astrocyte endfeet (A, cyan). Scale bar represents 1 µm (left). (b) TEM images of two blood vessels in the ME, (i) and (ii). Pseudocolors highlight different cells: cEC (purple), pericyte (teal), fibroblast (red), lumen (L, white) and neuropil (yellow). Insets show capillary fenestrations (white arrowheads), cEC tight junctions (white arrows), extracellular matrix-filled perivascular space (ECM), pericyte cells (P, teal) and fibroblast cells (F, red). Scale bar represents 1 µm. (c) TEM images of two groups of ME blood vessels. Scale bar represents 4 µm. (d) Representative images of cortex (i) vessel 1 and (ii) vessel 2 reconstruction by serial TEM. Pseudocolors show reconstructed regions: blood vessel lumen (L, red); EC (green); and pericyte cell (blue). White arrows indicate EC tight junctions (purple), and white boxes highlight ‘peg and socket’ pericyte-EC interactions. Scale bar represents 1 µm.

**Extended Data Fig. 9.**
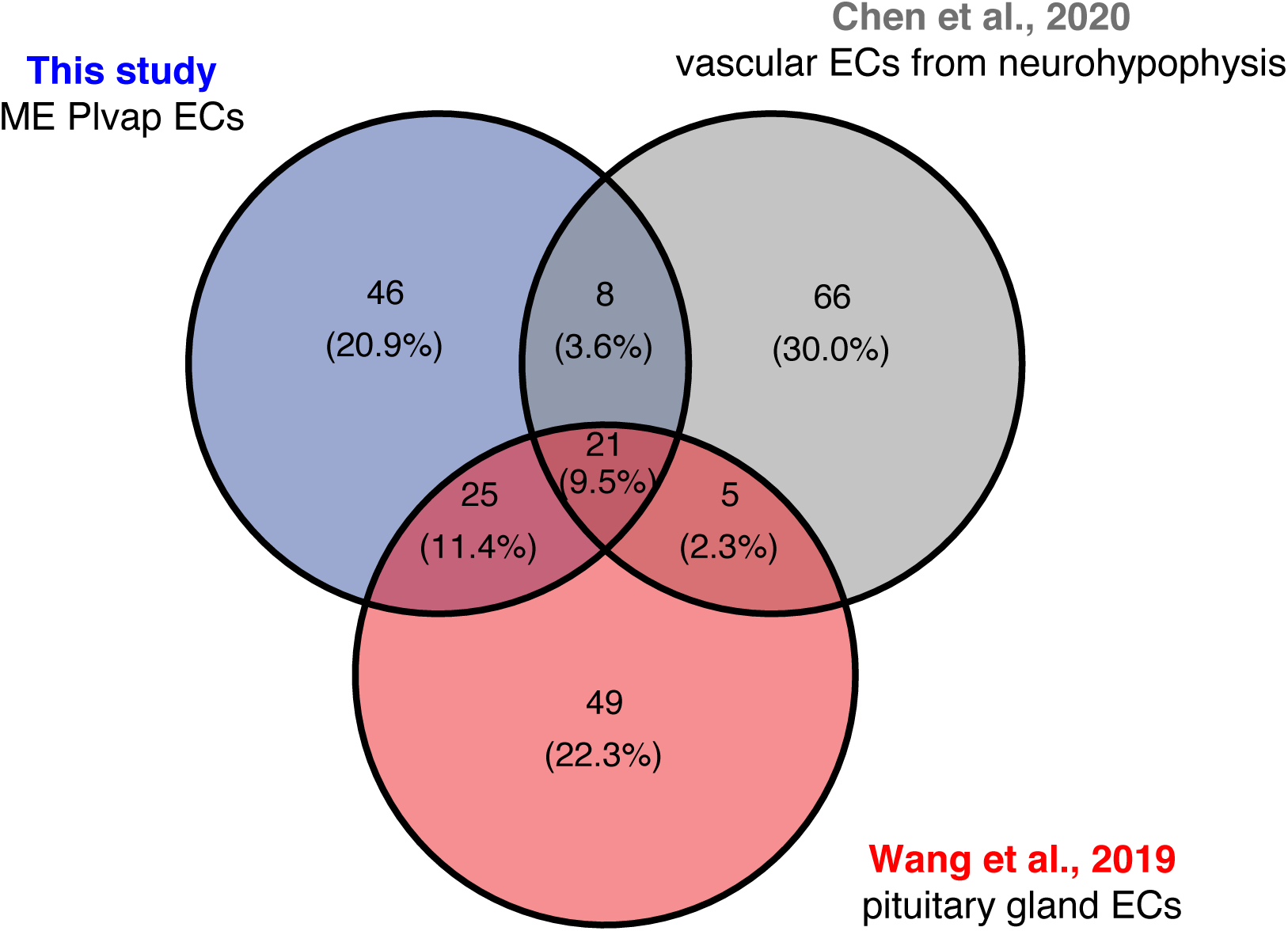
Comparison of scRNAseq of ME-derived Plvap+ ECs to published ECs from the mouse neurohypophysis and pituitary gland. The top 100 genes enriched in ME-derived Plvap+ ECs when compared to cortex-derived cECs (from Table 1; blue) were compared to the top 100 genes enriched in vascular endothelial cells from the mouse neurohypophysis^55^ (gray) and the top 100 genes enriched in ECs isolated from the mouse pituitary gland^40^ (red).

## Notes

### Competing Interest Statement

The authors have declared no competing interest.

